# Exploring the Relationship Between Extracellular Vesicles, the Dendritic Cell Immunoreceptor and MicroRNA-155 in an In Vivo Model of HIV-1 Infection to Understand the Disease and Develop New Treatments

**DOI:** 10.1101/2024.10.18.619157

**Authors:** Julien Boucher, Gabriel Pépin, Benjamin Goyer, Audrey Hubert, Wilfried Wenceslas Bazié, Julien Vitry, Frédéric Barabé, Caroline Gilbert

## Abstract

HIV-1 infection induces persistent immune system activation despite antiretroviral therapy. New immunomodulatory targets might be required to restore immune competence. The dendritic cells immunoreceptor (DCIR) can bind HIV-1 and regulate immune functions and extracellular vesicles (EVs) production. EVs have emerged as biomarkers and a non-invasive tool to monitor HIV-1 progression. In people living with HIV-1, an increase in the size and abundance of EVs is associated with a decline in the CD4/CD8 T cells ratio, a key marker of immune dysfunction. Analysis of host nucleic acids within EVs has revealed an enrichment of microRNA-155 (miR-155) during HIV-1 infection. Experiments have demonstrated that miR-155-rich EVs enhance HIV-1 infection in vitro. A humanized NSG-mice model was established to assess the *in vivo* impact of miR-155-rich EVs. Co-production of virus with miR-155-rich EVs heightened the viral load and lowered the CD4/CD8 ratio in the mice. Upon euthanasia, EVs were isolated from plasma for size and quantity assessment. Consistent with findings in individuals with HIV-1, increased EVs size and abundance were inversely correlated with the CD4/CD8 ratio. Next, by using the more closely related physiological virus co-product with EV-miR-155, we tested a DCIR inhibitor to limit infection and immune damage in a humanized mouse model. DCIR inhibition reduced infection and partially restored immune functions. Finally, viral particles and various EV subtypes can convey HIV-1 RNA. HIV-1 RNA was predominantly associated with large EVs (200-1000nm) rather than small EVs (50-200nm). Viral loads in large EVs strongly correlated with blood and tissue markers of immune activation. The humanized mice model has proven its applicability to studying the roles of EVs on HIV-1 infection and investigating the impact of DCIR inhibition.

**Author Summary:** Despite more than 40 years of research, HIV remains a threat to public health around the world. People living with HIV are efficiently treated with antiretroviral therapy, but damage to the immune system persists and the causes remain unknown. Extracellular vesicles allow material, such as microRNA, to transfer between cells. Here, we evaluated the impact of one microRNA, microRNA-155, transported by extracellular vesicles, on HIV infection. Mice were grafted with a human immune system to allow infection by HIV. We showed that extracellular vesicles carrying microRNA-155 amplified mice infection. Extracellular vesicles also reflect the state of their cell of origin. Their analysis can reveal biomarkers to monitor HIV infection. Thus, HIV viral load was quantified in purified extracellular vesicles. We found that the measurement of HIV viral load in purified EVs is a more precise biomarker of disease progression than the traditional plasma viral load. Additionally, potential treatments like DCIR inhibitors improve our ability to manage HIV-1 by restoring the CD4/CD8 ratio, a critical element of the infection process. Overall, our study highlighted the importance of extracellular vesicle cargo in a humanized mouse model of HIV-1 infection, as well as the potential of targeting DCIR to restore the immune response.

**Highlights:**

- MicroRNA-155 promotes HIV-1 infection of humanized NSG mice
- Abundance and size of total plasmatic EVs are biomarkers of immune dysfunction associated with HIV-1 infection
- DCIR inhibition limits HIV-1 infection of humanized NSG mice and attenuates immune impairment
- HIV-1 RNA enrichment in large EVs was associated with biomarkers of immune activation and dysfunction

## 1. Introduction

Antiretroviral therapy (ART) effectively reduces viral load in the periphery, but it cannot eradicate the virus, which remains primarily integrated into the genome of reservoir cells (1). Comorbidities associated with infection persist (2–5), possibly because sustained inflammation disrupts immune system responses (6, 7). The number of CD4 T lymphocytes (CD4TL) rebounds with treatment to an acceptable level (above 500 cells/µL), but the count of CD8 T lymphocytes (CD8TL) remains high (more than 500 cells/µL) (8, 9). CD8TL in HIV-1 patients exhibits differentiation defects (8, 10), a phenotype of exhaustion (8, 11, 12) and cellular activation (8, 13). Consequently, they eliminate infected cells less effectively (14, 15). Even if HIV-1 infection is successfully treated with ART (i.e., undetectable viral load and CD4TL count close to 500 cells/µL), circulating immune cells (CD8TL or myeloid-derived suppressor cells) still express varying degrees of exhaustion or immune response suppression phenotypes (16–19). These phenotypes are associated with disease progression or the development of non-AIDS-related comorbidities (20). They can be considered biological stigmas that are not restored by ART and affect immune competence. Identifying the underlying factors driving these dysregulations is imperative to elucidate the persistent cellular phenotypes. Moreover, uncovering novel biomarkers for monitoring immune activation and identifying new immunomodulatory targets is crucial.

One of the immune system modulators of interest in HIV-1 infection is the Dendritic Cell Immunoreceptor (DCIR) (21). DCIR, a 237-amino acid protein encoded by the human CLEC-4A gene, plays a pivotal role in immune homeostasis maintenance. Its significance extends across various domains, with documented involvement in autoimmune (22–25), inflammatory (25, 26) and infectious diseases (27–34). DCIR is constitutively expressed on myeloid cells and downregulated in an inflammatory context (35). In dendritic cells (DCs), DCIR internalization inhibits responses mediated by TLR8 and TLR9 (36, 37), two receptors known to play a role in innate immunity to viruses. DCIR signalling can also inhibit DC-SIGN-mediated antigen presentation (38), while DCIR inhibition blocks the production of Baff (39), a B-cell stimulatory factor. These negative immunoregulatory roles of DCIR in innate and acquired immunity make this lectin a relevant target for enhancing immune responses, as shown for bladder cancer treatment (40). Myeloid C-type lectin (CTL) receptors, a distinguished subset of pattern recognition receptors (PRRs), are pivotal in initiating immune responses. CTLs can recognize a broad spectrum of glycans, including the HIV-1 surface glycoprotein gp120, a highly glycosylated protein (41). The first step of HIV-1 infection involves the interaction of gp120 with CTLs, including DCIR (38, 42, 43). Then, virion-carrying DCs migrate to lymph nodes and other lymphoid tissues. DCIR plays a crucial role in mediating viral entry into the body by facilitating the transfer of virions to CD4TL (31) and releasing extracellular vesicles (EVs) (28).

EVs shuttle proteins and genetic material, including DNA, RNA, microRNAs, and other non-coding RNAs, which neighbouring cells can uptake (44). Functioning as intercellular messengers, EVs modulate the activities of recipient cells upon internalization of their cargo (44). These EVs are integral to normal physiological (45, 46) and pathological processes (47, 48). EVs include microvesicles (MVs) (100-1000 nm), exosomes (50-100 nm), apoptotic bodies (200-1000 nm), and virions (either fully infectious or not) (44, 49). In vitro studies have shown that EVs directly influence HIV infection (50–53) by maintaining (54) or activating viral reservoirs (55). EVs were found to transport viral components during HIV-1 infection. (56–58) The entire HIV RNA can be contained within EVs (59), and the RNA-binding element, TAR, can activate transcription and render naive CD4TL permissive to infection (51). Furthermore, the binding of HIV-1 to DCs leads to an increase in proapoptotic EV release (60). Using a DCIR inhibitor decreases the quantity of EVs released by DCs when stimulated by HIV-1 (28). EVs produced by CD4TL or DCs in contact with HIV can alter the responses of other immune cells to the virus and influence disease outcomes (50, 55, 56, 58, 61, 62).

HIV-1 infection or even contact with HIV-1 leads to changes in the expression profiles of microRNAs in CD4TLor macrophages (63), and EVs can contain this genetic material (64, 65). These microRNAs associate with specific mRNA sequences to inhibit their translation into proteins, thus influencing and modifying numerous cellular functions (66). Described in the modulation of different immune responses and highly expressed in PLWH (63, 67, 68), cellular miR-155 is mainly found in lymphoid organs, particularly the spleen and thymus (69). Peripheral blood mononuclear cells (PBMCs) of PLWH express miR-155 at a higher level than non-infected subjects (70). Cellular overexpression of miR-155 is associated with a lower CD4/CD8 ratio, immune activation and exhaustion (70). Overexpression of miR-155 in PLWH CD8TL correlated with CD8TL activation and differentiation (71). Our previous studies showed that HIV preparations and inflammatory molecules increase miR-155 expression in peripheral blood mononuclear cells (PBMCs), that miR-155 is predominantly enriched in large EVs, and that EV-miR-155 amplified HIV-1 replication by promoting an inflammatory environment (50).

EVs are produced in tissues during infection, circulate in biological fluids and are relatively reliable biomarkers of tissue activation status, particularly concerning immune system cells (72–74). Mainly through their protein or microRNA contents, they are considered liquid biopsies in oncology (75, 76), and this concept could soon expand to the field of infectious diseases. The content of EVs in microRNA has been little studied in the context of HIV infection (77). Moreover, many studies focus on a single population of EVs, either small EVs (exosomes) or large EVs (MVs), resulting in limited information obtained from the same samples on different types of EVs and even less on apoptotic EVs. We discovered in the plasma of PLWH that the abundance and size of EVs correlated with the number of CD8TL (50, 64, 78, 79). We have developed robust and quantitative methods to measure microRNAs in the two populations of EVs (50, 78) and to express their quantity in copies per EV (79–83). We showed that the number of EVs and the quantification of miR-155 per small EVs predict viral rebound under antiretroviral therapy (ART) (80). Furthermore, the measurement of miR-155 in large EVs predicts a CD8TL count above 500 cells/µL (79). These measurements of miR-155 quantity per EVs are significant for predicting immune activation and viral rebound (79, 80), and microRNAs in EVs may act as epigenetic factors (84, 85).

EVs appear to be a cornerstone of immune activation triggered by HIV-1 infection. Understanding, monitoring, and ultimately managing this ongoing process is crucial for improving the long-term health outcomes of PLWH. However, no study has examined the role of miR-155-enriched EVs in immune activation in a mouse model of HIV-1 infection, nor has the importance of DCIR in this process as well as the value of plasmatic EVs as biomarkers. The most used mouse strains for HIV-1 infection are NOD.Cg-*Prkdc^scid^Il2rg^tm1Wjl^*/SzJ (NSG) mice and NOD.Cg-*Prkdc^scid^Il2rg^tm1Sug^*/ShiJic (NOG) mice. (86, 87). Both mouse models have a deficiency in IL-2rg, resulting in the absence of several immune cells. HuCD34 NSG mice can be infected with HIV-1 through various routes, including intraperitoneal, vaginal, or rectal routes (86). These mice have been used to assess responses to antiretroviral treatment or to monitor HIV pathogenesis (86, 88). Using miR-155-enriched EV preparations and DCIR inhibitors (29), we have established that huCD34 NSG mice are suitable for studying EVs functional roles and biomarkers of EVs. Specifically, this study highlights that viral RNAs present in large EVs are biomarkers of immune activation and that DCIR appears to be a relevant target for restoring immune response.

## 2. Results

### 2.1 The roles of EVs during HIV-1 infection of humanized mice

EV-miR-155 are rapidly released by immune cells during HIV-1 infection and amplifies HIV-1 infection in *in vitro* settings (50). To answer this question, we chose to use viruses produced by transfection in HEK-293T cells, since these cells are negative for miR-155 and the majority of infection studies use viral productions from these cells. Here, we tested their effect in an *in vivo* model of HIV-1 infection with humanized NSG mice. Mice were inoculated with the equivalent of 5000 TCID50 of HIV-1 co-produced with miR-155-rich EVs (EV-miR-155) or not (EV-Mock) (Fig 1A and S1A-B). Enrichment of miR-155 in EVs was confirmed by qPCR (Fig S1C). The concentrations of miR-92 and miR-223 were measured to ensure that their concentration remained constant between the two viral preparations (Fig S1C). In addition, co-produced virus and EVs were purified on an iodixanol velocity gradient to separate EVs and viruses (89). MiR-155 quantification in the gradient fractions showed miR-155 in EVs fractions and virus fractions (Fig S1D). Mice were weighed weekly to ensure that they maintained a healthy weight (Fig 1B). Disease progression was monitored by the CD4/CD8 ratio measurement. At 14 days post-infection, infected mice that received virus co-produced with EV-miR-155 had the lowest CD4/CD8 ratio. This tendency was maintained throughout the experiment (Fig 1C).

**Figure 1.**
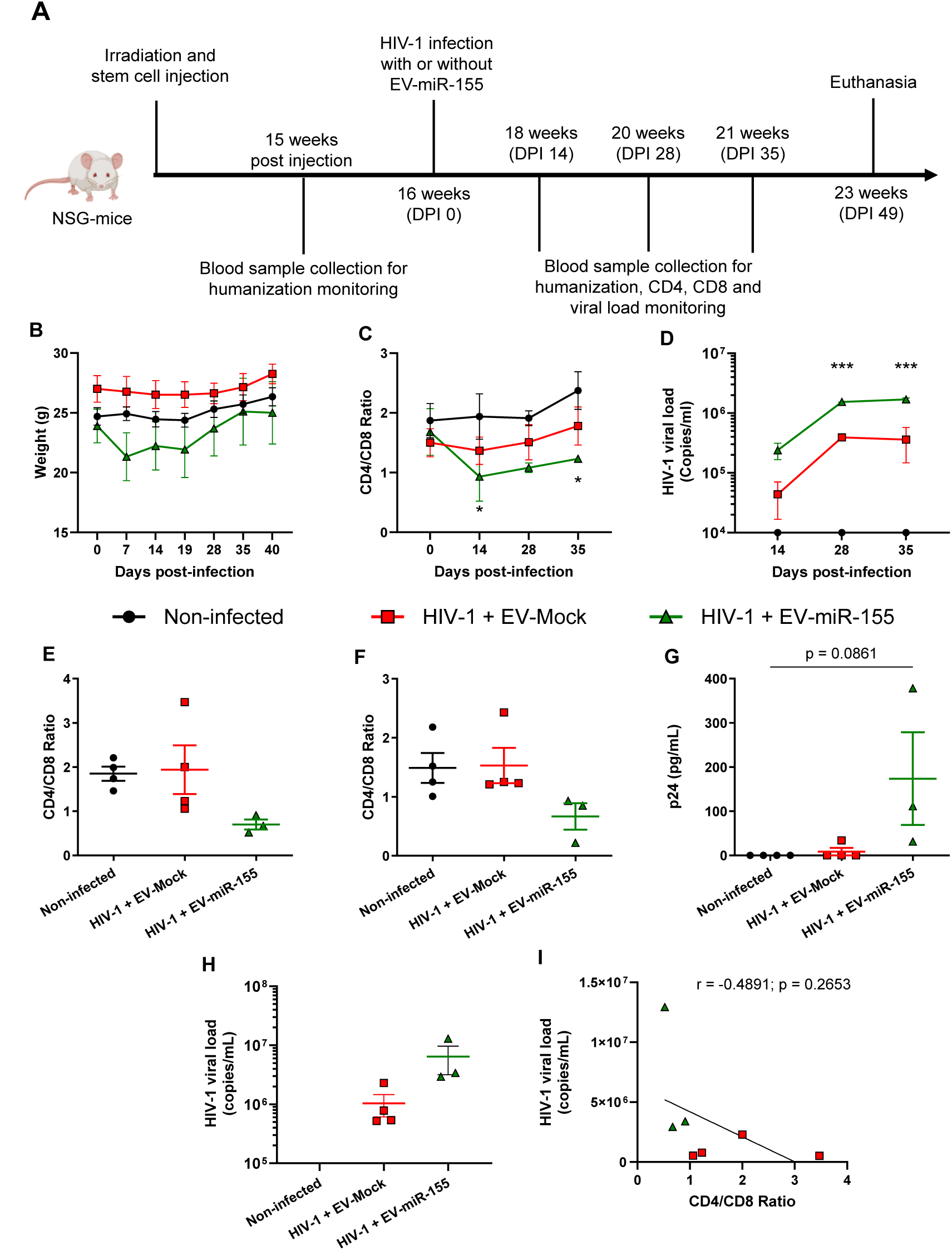
MiR-155-rich EVs promote HIV-1 infection in humanized mice. Eleven humanized NSG mice were used for the protocol: Four were inoculated with sterile PBS (control group), four with HIV-1 + EV-Mock (130 ng p24; 5000 of TCID 50; NL4.3Balenv+ EV-Mock group) and three with HIV-1+ EV-miR-155 (90ng p24, 5000 of TCID 50; NL4.3Balenv + EV-miR-155 group) **A.** Timeline of mice experiments **B.** Mice’s weight was monitored weekly. **C.** Immune cells were purified from the blood and the CD4/CD8 ratio was monitored by flow cytometry at 14, 28 and 35 DPI. Two-way ANOVA was used for statistical analysis of CD4/CD8 ratio with the control group as the reference (* p < 0.05). **D.** Plasma viral load was measured by RT-qPCR with a specific primer set targeting the virus LTR region. Two-way ANOVA was used for statistical analysis of the plasmatic viral load with the HIV-1 group as the reference (*** p < 0.001). **E and F**. CD4/CD8 ratio at euthanasia in blood and spleen. **G**. Viral particles were detected in plasma at euthanasia using anti-p24 ELISA **H**. Plasma viral load at euthanasia. A one-way ANOVA statistic test was performed for statistical analysis. **I.** Correlation between plasma viral load and blood CD4/CD8 ratio at euthanasia for all infected mice. DPI: days post-infection. The mouse image was obtained via BioRender.

To ensure reliable detection of HIV-1 RNA in mice plasma, cDNA pre-amplification was performed to increase the number of template copies, allowing easier detection of HIV-1 RNA by RT-qPCR (90) (Fig S2 and Table S1). As expected, a higher viral load was measured in mice that received EV-miR-155 at all time points (Fig 1D). At euthanasia, mice that were infected in the presence of EV-miR-155 had a lower blood (Fig 1E) and spleen (Fig 1F) CD4/CD8 ratio than the controls. Mice infected in the presence of EV-miR-155 had the highest concentration of capsid protein p24 (Fig 1G) and viral RNA (Fig 1H). Consequently, a higher viral load was associated with a lower CD4/CD8 ratio (Fig 1I).

Expression of HLA-DR, a biomarker of immune activation (91), and PD-1, a biomarker of immune exhaustion (92), was measured on CD4TL and CD8TL (Fig S3A). HLA-DR expression was increased on CD8TL, and PD-1 was overexpressed on both CD4TL and CD8TL from mice exposed to HIV-1 + EV-miR-155 (Fig S3B). CD8TL were purified from the spleen and reactivated for nine days to assess their capacity to respond to TCR activation and to validate immune exhaustion. Both at resting and reactivated states, EV-miR-155 was associated with increased expression of PD-1 on CD8TL (Fig S4). These results show that EV-miR-155 increased mice infection and potentialized immune dysfunction.

At euthanasia, total EVs were purified from plasma to characterize EVs as biomarkers during HIV-1 infection of humanized mice. The size and abundance of EVs were analyzed, as both parameters are known to inversely correlate with the CD4/CD8 ratio (64). EV hydrodynamic size measurement revealed that infection in the presence of miR-155 increased EV size (Fig 2A). EV abundance was estimated with the measure of the AChE activity. Infection in the presence of miR-155 was associated with a notable increase in total plasma EV abundance (Fig 2B). As expected, total plasmatic EV size (Fig 2C) and abundance (Fig 2D) negatively correlated with the CD4/CD8 ratio in HIV-1 infected mice. EV size and abundance also correlated with the plasma viral load at euthanasia (Fig 2E and F). These results show that the humanized mouse model is suitable for studying the role of EVs as biomarkers during HIV-1 infection. Therefore, EV-miR-155 were co-produced with viral stocks in HEK293T to be closer to physiological conditions as we have previously shown (50, 81). Finally, these results highlight the importance of this microRNA mainly produced by immune cells in HIV-1 pathogenesis. Viruses co-produced with EV-miR155 were used to pursue our understanding of the mechanisms of immune activation, the determination of new biomarkers and the characteristation of new targets to regulate immune activation,..

**Figure 2.**
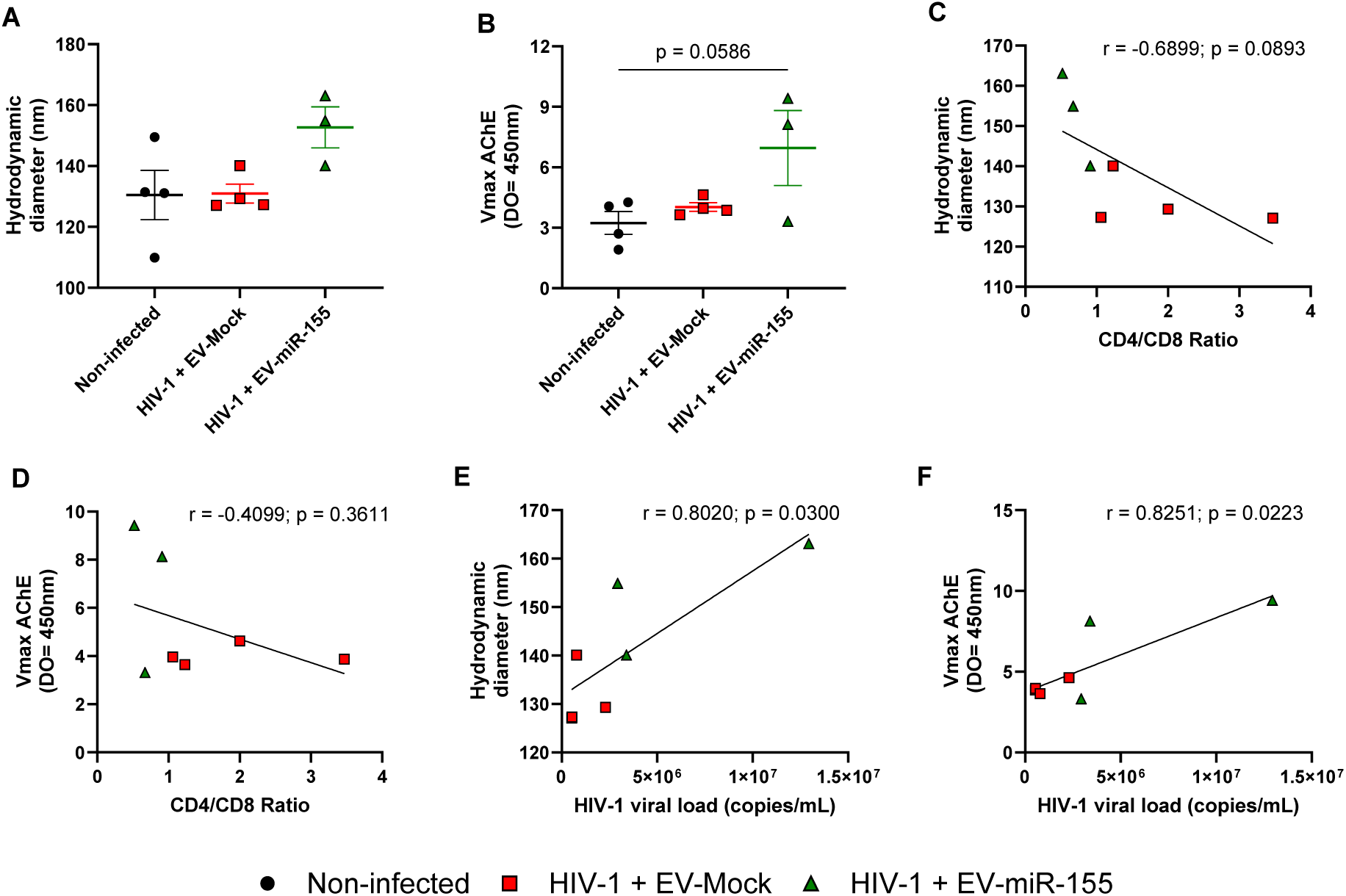
EVs size and abundance correlate with a lower CD4/CD8 ratio. Total EVs from the plasma of mice were purified using ExoQuick. **A**. The hydrodynamic diameter of vesicles from plasma was estimated using Nanosizer. **B**. AChE activity (Vmax, D.O. 450nm/min) was measured using an enzymatic assay. One-way ANOVA was used for statistical analysis. **C.** Correlation between plasmatic EV size and blood CD4/CD8 ratio. **D.** Correlation between AChE activity and blood CD4/CD8 ratio. **E.** Correlation between plasmatic EV size and plasma viral load. **F.** Correlation between AChE activity and plasma viral load.

### 2.2 The effect of DCIR inhibition on HIV-1 infection and immune dysfunction in humanized mice

Blockade of DCIR with an inhibitor (1-benzofuran-2-yl(phenyl)methanone) prevents HIV-1 binding to the cell (29). The DCIR inhibitor can also downregulate EV production (28). It has proven its *in vitro* efficacy to mitigate HIV-1 infection but has never been tested in *in vivo* settings. The impact of mouse treatment with a DCIR inhibitor before virus inoculation was evaluated. First, human DCIR expression was measured on the cells and tissues of the humanized mice. Mice were humanized with the injection of human hematopoietic stem cells. At 13 weeks post-injection and at multiple subsequent time points (Fig 3A), human engraftment was determined by the measure of human CD45+ cell proportions in mice blood (Fig 3B). Preliminary experiments by flow cytometry confirmed that the highest proportion of DCIR+ immune cells was in the spleen. DCIR+ cells were also found in the bone marrow, and few were present in the blood of mice (Fig S5). In both the spleen and the bone marrow, DCIR+ cells were mostly of myeloid origin (Fig S5B-E). In the spleen, most myeloid cells were CD4+ DCIR+ (Fig S5F), in the bone marrow, most myeloid cells were CD4+ DCIR-(Fig S5G).

**Figure 3.**
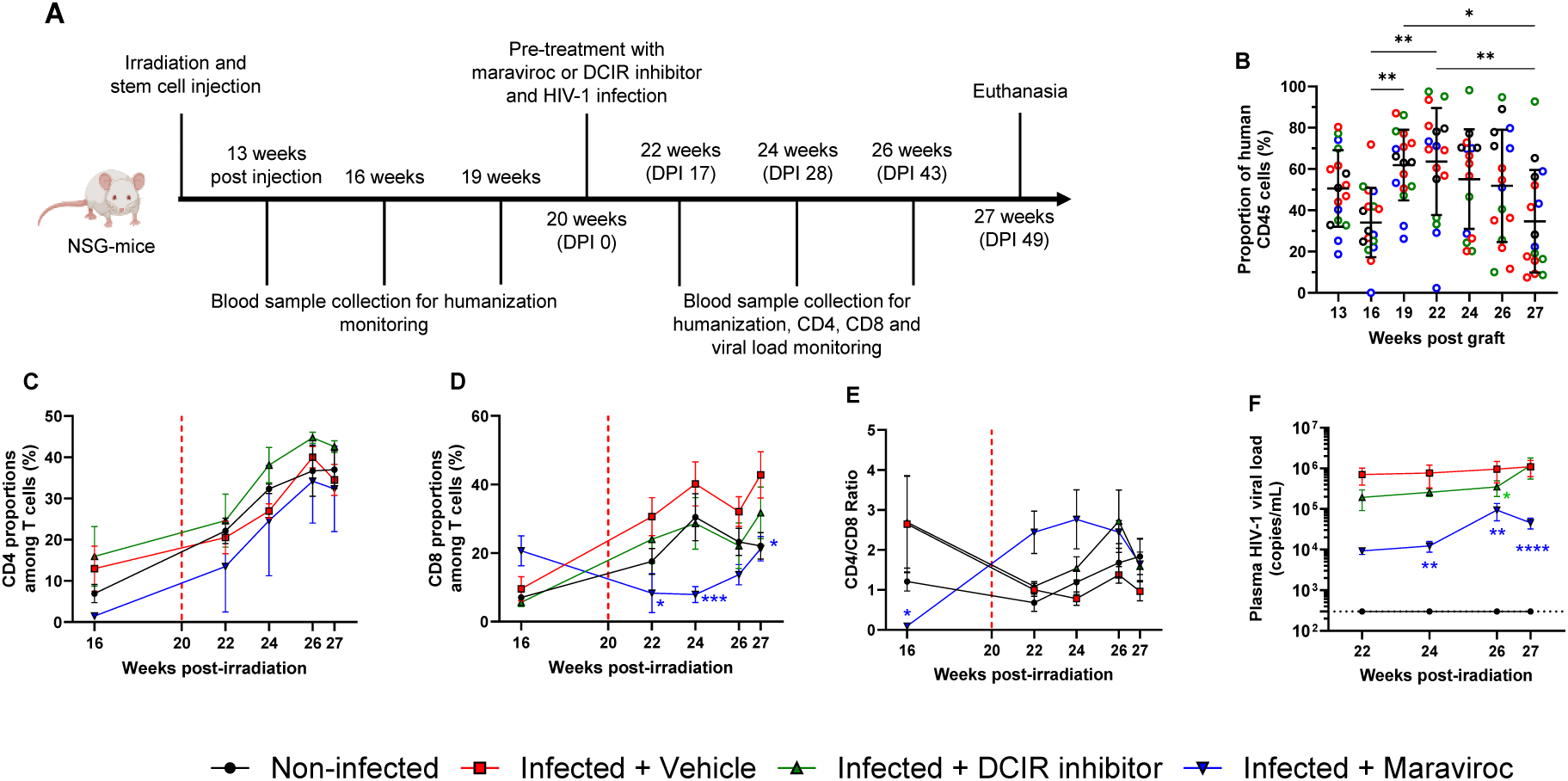
The effect of DCIR inhibitor or maraviroc pre-treatment on humanized NSG mice infection. Humanized NSG mice were infected with the NL4.3Balenv + EV-miR-155 viral preparations. Three groups of mice received the viral preparation: an infected + hydroxyethylcellulose group as the group of reference (n = 7), and two groups were inoculated before infection with DCIR inhibitor (n = 4) or maraviroc (n = 3). Non-infected mice were controls (n=4). **A.** Timeline of mice experiments (DPI: days post-infection). **B.** Engraftment of hematopoietic stem cells in mice was measured in blood samples. The proportion of human CD45+ cells was determined by flow cytometry. **C.** Proportion of CD4+ cells among T cells were assessed by flow cytometry. **D.** The proportion of CD8+ cells among T cells was assessed by flow cytometry. **E.** Peripheral blood CD4/CD8 ratio after infection calculated with flow cytometry data. **F.** Plasma viral load was measured at various time points. Data presented are mean with standard error of the mean (SEM). Statistical analysis was carried out by two-way ANOVA to compare the different mice subgroups (* p < 0.05; ** p < 0.01; *** p < 0.001; **** p < 0.0001). The mouse image was obtained via BioRender.

The results described above shown that virus produced on HEK-293T enriched in EV-miR155 are more relevant to infect mice. A group of non-infected mice was the control group, and the remaining mice were infected with viral preparation enriched in EVs bearing miR-155., Mice that received NL4.3Balenv + EV-miR-155 received 7.31×10^6^ total copies of EV-borne miR-155. This is a physiological concentration because we previously observed that 10^6^ PBMCs released around 10^6^ copies of EV-borne miR-155 in 24 hours (50). Among infected mice, some were pre-treated with maraviroc to attenuate mice infection (93) and act as a control, and others were pre-treated with a DCIR inhibitor. For the pre-treatment, the DCIR inhibitor (10 mM) or maraviroc (5 mM) diluted in PBS/2.2 % hydroxyethylcellulose gel was intra-vaginally applied 60 minutes before infection.

CD4TL and CD8TL were monitored by flow cytometry. CD4TL proportions were similar between all groups, except mice that received DCIR with a slightly higher proportion (Fig 3C). CD8TL proportions increase in infected mice compared to control mice. DCIR inhibitor moderately diminished CD8TL counts, and maraviroc significantly reduced CD8TL counts (Fig 3D). CD4/CD8 ratio was lower in infected mice compared to control mice. The CD4/CD8 ratio was restored in maraviroc-treated mice at 28 days post-infection and in DCIR inhibitor-treated mice at 43 days post-infection (Fig 3E). Finally, plasma viral load was measured at 17, 28, 43 and 49 days post-infection. As shown in Figure 3 and highlighted in Supplementary Figure 6, the DCIR inhibitor group had a slightly lower viral load than the virus-only group. Maraviroc had a greater effect in inhibiting the infection of mice (Fig 3F).

Viral load was also measured in immune cells purified from the spleen and the lungs. As observed in the plasma, non-treated mice had the highest viral load (Fig 4). Mice pre-treated with the DCIR inhibitor had a similar viral load in the spleen (Fig 4A) and a slightly lower viral load lungs (Fig 4B). Viral load was undetectable in the spleen of all three mice and in the lungs of one out of three mice in the maraviroc group (Fig 4).

**Figure 4.**
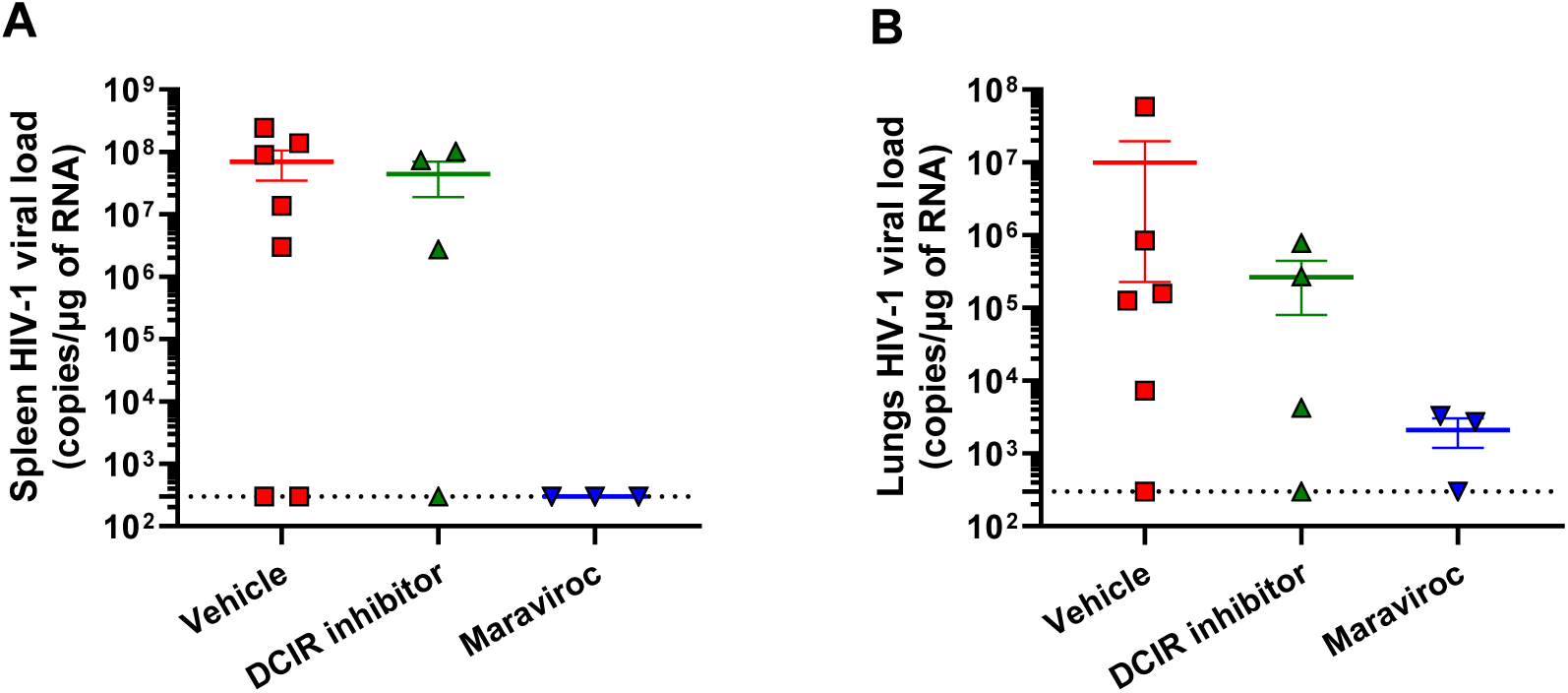
Viral load measurement in spleen and lungs immune cells. Spleens and lungs were processed with a gentleMACS dissociator. Tissues were deposited on 40µm-pore size nylon filter to collect cells. Lung cells were further purified on Cytvia Percoll centrifugation medium. Finally, erythrocytes were lysed with an ammonium chloride solution. **A.** Viral load measurement in the spleen. **B.** Viral load measurement in the lungs. Statistical analysis was carried out by one-way ANOVA.

Immune dysfunction in mice was further evaluated with purified immune cells from the spleen, the lungs and the liver. Analysis of human CD45 expression showed that human leucocyte amount in the spleen, the liver and the lungs are unaffected by infection and treatment (Fig S7A-E-I). In the spleen, no notable variation of CD4TL proportions was observed (Fig S7B). Spleen CD8TL proportions increased in infected mice, but not in maraviroc-treated mice (Fig S7C). The CD4/CD8 ratio was the lowest in infected mice and restored in mice treated with DCIR inhibitor or maraviroc (Fig S7D). Similar results were observed in the liver (Fig S7F-H). In the lungs, CD4TL, CD8TL and CD4/CD8 ratio variations were negligible (Fig S7J-L).

In addition, HLA-DR and PD-1 expression by blood and spleen CD8TL were measured. For this analysis, CD8TL were divided into low CD8 expression (CD8^low^) and high CD8 (CD8^high^) expression. Repeated antigen stimulation increases the proportion of activated CD8^low^ TL (94), which is common in PLWH (95). A higher proportion of CD8^low^ TL was measured in the blood and the spleen of infected mice than in control mice (Fig S8). DCIR inhibitor and maraviroc-treated infected mice blood CD8^low^ TL proportions were reduced to the level of control mice (Fig S8). In the spleen, CD8^low^ TL were back to the control level in DCIR inhibitor-treated mice (Fig S8). In infected mice, HLA-DR and PD-1 expression is increased by blood and spleen CD8^low^, CD8^high^, and CD8TL, which was mitigated by maraviroc and DCIR inhibitor treatments (Fig S9). The effect of DCIR inhibition was greater in CD8^low^ TL, while the effect of maraviroc was stronger towards CD8^high^ TL (Fig S9).

### 2.3 Purified EVs viral load correlates with blood biomarkers of immune impairment and HIV-1 spreading in mouse tissues

Large and small EVs were purified from the plasma collected at euthanasia. Purified EV subtypes were quantified by flow cytometry as described in the material and methods (Fig S10). The hydrodynamic size of large and small EVs was measured by dynamic light scattering with a Nanoszier instrument. Large EVs were bigger than small EVs (Fig S11A-B). Relative quantification by dynamic light scattering and absolute quantification by flow cytometry show that small EVs are increasingly more abundant than large EVs over time (Fig S11C-D). Exosomes (small EVs) resist treatment with SDS 0.025%, while microvesicles (large EVs do not (96). Both subtypes were treated with SDS 0.025% or SDS 0.125% before cytometer acquisition. Most large EVs were lysed by SDS 0.025% (Fig S11E). Small EVs resisted the treatment with SDS 0.025% but not with SDS 0.125% (Fig S11F). Small EVs were more abundant than large EVs (Fig S11). Infected mice and infected mice treated with DCIR inhibitor had a lower large EVs concentration than the control mice (Fig S12A). Inversely, infected mice treated with maraviroc had a higher large EVs concentration (Figure S12A). Absolute small EVs counts were stable among all mice subgroups (Fig S12B). In the plasma, only maraviroc-treated mice had a significantly lower plasma viral load (Fig 5A). In purified plasmatic large EVs, viral load was diminished in DCIR inhibitor and maraviroc-treated mice (Fig 5B-D). No change was observed in small EVs (Fig 5C-E). Total plasma contained more viral RNA than both EV subtypes and large EVs contained significantly more HIV-1 RNA than small EVs (Fig S13).

**Figure 5.**
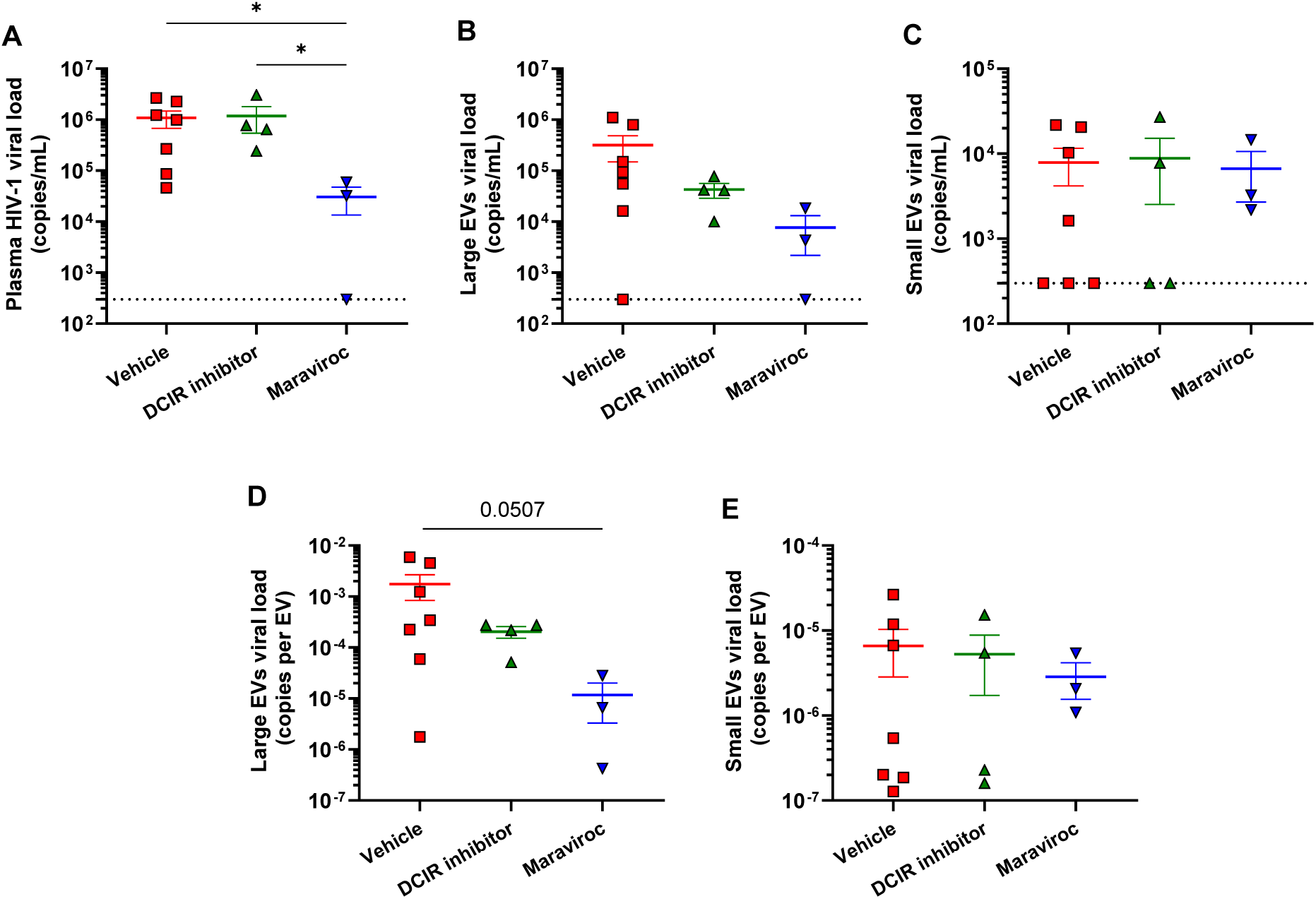
Viral load distribution in plasmatic EVs subtypes in humanized mice. **A.** Plasma viral load was measured at euthanasia. Large and small EVs were purified from the plasma of mice at euthanasia. Viral load was measured in each EV subtype by RT-qPCR. **B.** Large EVs viral load for mice subgroups. **C.** Small EVs viral load for mice subgroups. **D.** The viral load value, in terms of copies per mL, was divided by the concentration of EVs per mL to obtain the number of HIV-1 RNA copies per EV. The number of HIV-1 RNA copies per large EV. **E.** The number of HIV-1 RNA copies per small EV. Statistical analysis was carried out by one-way ANOVA (* p < 0.05).

The correlations between plasmatic EVs viral load and known biomarkers of immune dysfunction (CD4TL count, CD8TL count and CD4/CD8 ratio) associated with HIV-1 infection were calculated (Table 1). Plasma viral load did not correlate with the CD4TL count, the CD8TL count or the CD4/CD8 ratio. We found correlations between large EVs viral loads, higher CD8TL counts and a lower CD4/CD8 ratio. Small EVs viral load correlated with none of the three biomarkers.

**Table 1.** Correlation of mice plasma EVs viral load with biomarkers of HIV-1 pathogenesis.

|  | CD4+ T cell count | CD8+ T cells count | Blood CD4/CD8 ratio |
| --- | --- | --- | --- |
| Plasma viral load (copies/mL) | r = -0.1638; p = 0.5758 | r = 0.3948; p = 0.1625 | r = -0.2797; p = 0.3328 |
| Large EVs viral load (copies/mL) | r = -0.2002; p = 0.4892 | **r = 0.6491; p = 0.0141** | r = -0.5017; p = 0.0698 |
| Large EVs viral load (copies/EV) | r = -0.2141; p = 0.4623 | **r = 0.7982; p = 0.0006** | r = -0.4770; p = 0.0846 |
| Small EVs viral load (copies/mL) | r = 0.2214; p = 0.4469 | r = -0.2328; p = 0.4232 | r = 0.3534; p = 0.2152 |
| Small EVs viral load (copies/EV) | r = 0.2442; p = 0.4002 | r = -0.0778; p = 0.7914 | r = 0.1673; p = 0.5676 |

We calculated the correlation between the plasma viral load or the purified EVs viral load and the virus spreading in the mice tissues. Plasma and small EVs viral load did not correlate with the spleen nor the lungs viral load. On the other hand, the large EVs viral load correlated strongly with the spleen viral load at euthanasia and reactivated spleen cells viral load (Table 2). In addition, the large EVs viral load correlated positively with the CD8TL counts in the spleen (Table S2). No correlation was found between EVs viral load and CD4TL, CD8TL or CD4/CD8 ratio in the lungs (Table S3) and the liver (Table S4).

**Table 2.**
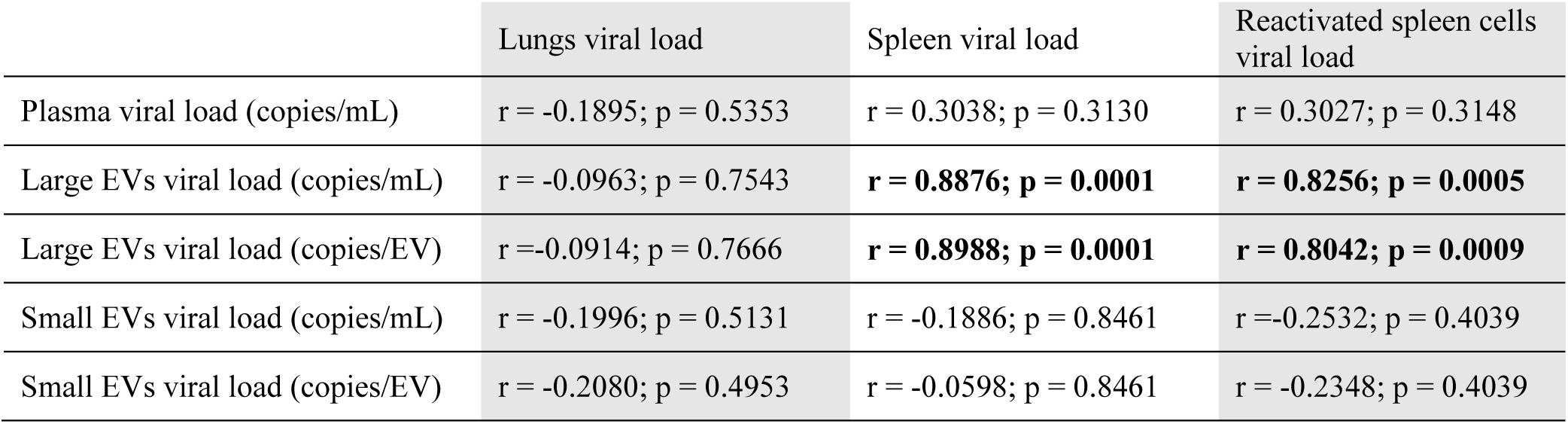
Correlation of mice plasma EVs viral load with spleen, lungs and reactivated spleen cells viral load.

|  | Lungs viral load | Spleen viral load | Reactivated spleen cells viral load |
| --- | --- | --- | --- |
| Plasma viral load (copies/mL) | r = -0.1895; p = 0.5353 | r = 0.3038; p = 0.3130 | r = 0.3027; p = 0.3148 |
| Large EVs viral load (copies/mL) | r = -0.0963; p = 0.7543 | **r = 0.8876; p = 0.0001** | **r = 0.8256; p = 0.0005** |
| Large EVs viral load (copies/EV) | r = -0.0914; p = 0.7666 | **r = 0.8988; p = 0.0001** | **r = 0.8042; p = 0.0009** |
| Small EVs viral load (copies/mL) | r = -0.1996; p = 0.5131 | r = -0.1886; p = 0.8461 | r = -0.2532; p = 0.4039 |
| Small EVs viral load (copies/EV) | r = -0.2080; p = 0.4953 | r = -0.0598; p = 0.8461 | r = -0.2348; p = 0.4039 |

Every correlation was also calculated with the sum of large and small EVs viral loads to determine if the total EVs viral load remained a biomarker of disease progression. Total EVs viral load maintained the same correlations as large EVs.

As shown in the supplementary Figure 4, the exhaustion phenotype seemed to be imprinted in the infected cells from the spleen. Purified spleen cells were reactivated with an anti-CD3/CD28 cocktail. We compared the expression of activation (HLA-DR) and exhaustion (PD-1) markers on T cells from non-treated mice and DCIR or maraviroc-treated mice (Fig S14). HLA-DR expression slightly diminished on CD4TL and CD8TL from DCIR inhibitor-treated mice. Maraviroc-treated mice had lower HLA-DR expression on CD4TL. As shown, PD-1 expression increased in both cell types in infected mice when compared to non-infected mice. Mice treated with DCIR inhibitor seemed to have the lowest PD-1 expression among all mice. (Fig S14).

## 3. Discussion

The multiple roles of EVs in HIV-1 infection are increasingly documented (97). EVs appear to be a cornerstone of immune activation triggered by HIV-1 infection. Understanding, monitoring, and ultimately managing this ongoing process is crucial for improving the long-term health outcomes of PLWH. This study aimed to develop and characterize an in vivo model to elucidate the mechanisms driving immune activation by introducing EVs containing miR-155 into viral preparations. Through analysis of viral components in plasma EVs, this study has addressed a crucial aspect of monitoring immune activation in humanized mice. Finally, the *in vivo* exploration of a DCIR inhibitor as a strategy to control immune activation has revealed promising results.

In the first mice experiment, the virus co-produced with EVs-miR-155 showed a priming effect on HIV-1 infection. Based on *in vitro* experiments, miR-155-rich EVs are rapidly released by immune cells in response to HIV-1 (50). Viral preparations produced by HEK293T cells contain EVs with low concentrations of miR-155. Thus, EVs bearing miR-155 render our viral preparations more closely related to viruses produced by immune cells. In humanized mice, miR-155-rich EVs increased viral replication, as previously observed in PBMCs (50). Mice infected by virus co-produced with miR-155 have shown a greater immune dysfunction, determined by a lower CD4/CD8 ratio in both blood and spleen and an exhausted phenotype on spleen T cells. More importantly, conserving PD-1 expression on the reactivated spleen CD8TL (Fig S3) strengthens the potential link between the epigenetic transformation of CD8TL by EVs bearing miR-155 as shown in human CD8TL (98). CD4/CD8 ratio and PD-1 expression on CD8TL are respectively biomarkers of immune dysfunction and exhaustion in PLWH (8, 92, 99). These results confirmed that EV-borne miR-155 influences HIV-1 infection in a dynamic more closely related to HIV pathogenesis observed on PLWH. In PLWH, miR-155 enrichment in EVs was identified as a biomarker of immune activation and viral replication (80, 83). Our *in vivo* results validate that miR-155 is more than a biomarker; it is directly involved in virus dissemination early in HIV-1 infection. Additionally, our findings suggest its potential involvement in the permanent modification of CD8TL exhaustion phenotypes as evidenced by the sustained elevation of PD-1 expression observed in reactivated cells from infected mice spleen across two independent experiments (Fig S3, S9).

Knight et al. initially demonstrated HIV-1 infection in dendritic cells (100). The involvement of DCIR in the productive infection of dendritic cellshas emerged more recently, as well as its role in proapoptotic EVs release (28). In the second mice experiment, the DCIR inhibitor was tested in humanized mice to limit HIV-1 infection and EV release. The results showed that treatment with DCIR inhibitor reduced mouse viral load to a lesser extent than maraviroc. Although the results observed are modest, they remain encouraging.

Further evidence that the DCIR inhibitor had an effect is shown in Figure S4H-I, where the number of DCIR-positive cells is increased in the spleen but not in the bone marrow. Crucially, DCIR is expressed on both infected and apoptotic CD4TL cells (32), and DCIR inhibitors decrease apoptosis induced by HIV-1 in CD4TL. These previous observations are aligned with the result in Figure 3C, where the CD4TL proportion is increased in DCIR inhibitor pre-treated mice. Blocking DCIR on infected CD4TL, in addition to preventing the initiation of the viral infection process by DC, could potentially enhance immune system responses already compromised by infection, restoring the CD4TL and limiting the proliferation and exhaustion of CD8TL (Fig 3). DCIR inhibition is known to reduce the number of EVs released by dendritic cells upon HIV-1 infection (28). Unfortunately, in our mice experiments, DCIR inhibition did not affect EV concentration in plasma. Further optimization of experimental parameters of DCIR inhibitor treatment is required to maximize its effects. In addition to treatment with the DCIR inhibitor less than 60 minutes to infection, we could increase the compound concentration or explore the possibility of intravenous injection post-infection in future experiments.

We also evaluated the reliability of humanized models to study the role of EVs as biomarkers of HIV-1 infection progression. Plasmatic EVs size and abundance by AChE activity measurement negatively correlated with the blood CD4/CD8 ratio in humanized mice, which corroborates previous observations in PLWH (64). Therefore, humanized NSG mice appear to be a suitable model to study plasma EVs as biomarkers to monitor HIV-1 infection. EVs and HIV-1 particles are purified in the same fractions when we use ExoQuick (101). Thus, EVs contain HIV-1 components (51, 102) and HIV-1 particles contain host molecules (103). To our knowledge, studies on HIV-1 RNA packaging in EVs are mostly focused on exosomes (referred to as small EVs in this study). This could be explained by the observations that HIV-1 proteins interact with host proteins involved in small EVs biogenesis mechanisms, which causes content interchange between virus and small EVs (104). One study showed that monocyte-derived macrophage microvesicles contained infectious HIV-1 particles (105).

The humanized NSG-mice model experiments confirmed that HIV-1 RNA is preferentially associated with large EVs. Maraviroc and DCIR inhibitors lowered total plasma and large EVs viral load and did not affect small EVs viral load. Several correlation analyses show that viral materials in large EVs were associated with a weak CD4/CD8 ratio in blood and tissues and a permanent exhaustion phenotype of CD8TL in the spleen than plasma viral load (Table S5). Therefore, it could be of greater interest to measure viral load in large EVs rather than small EVs since they can be concentrated by ultracentrifugation at 17,000 x *g* only. This could facilitate the detection of viral replication at very low levels.

We quantified significantly more viral RNA associated with large EVs than small EVs. However, we cannot conclude that functional HIV-1 particles are associated with or contained in large EVs, even if we measured HIV-1 RNA in large EVs. Treating mice with maraviroc lowered viral load in large EVs but did not affect small EVs. In other words, decreasing viral replication only decreased viral load in large EVs. This suggests that the newly produced virus was mostly associated with large EVs. This hypothesis is under investigation by an *in vitro* infectivity assay to determine the proportion of functional HIV-1 particles associated with large and small EVs, further purified by a velocity gradient. Such experiments require a large amount of EVs and viruses, which was limited by the small volume of mice plasma. MiR-155 can increase HIV-1 infectivity and is strongly enriched in large EVs (50). MiR-155 could potentiate the infectivity of large EV-associated HIV-1 particles. Neutrophils are a significant source of large EV-borne miR-155 (106). Although the humanized NSG mice model develops functional neutrophils, their numbers remain relatively low (107). This could explain the low concentration of miR-155 in plasmatic EVs of mice. Injection of cytokines, such as granulocyte-colony stimulation factor (G-CSF), has enhanced neutrophil counts in humanized mice and it could improve our mice model (108).

We found that the measurement of viral load in purified large EVs is a better indicator of immune dysfunction and virus dissemination in the blood, the spleen, and the liver than the standard plasma viral load measurement. Plasma viral load and the CD4TL count are routinely measured in PLWH as part of their clinic monitoring to confirm treatment efficacy and to detect viral rebound (109).

The CD8TL counts and the CD4/CD8 ratio are well-established predictors of disease progression and non-AIDS comorbidities (110). However, CD8TL count and CD4/CD8 ratio are rarely considered during clinical testing (110). CD4TL and CD8TL counts require more equipment, reagents, expertise, and time than viral load quantification. Our data showed that the distribution of viral RNA in purified EV subtypes changed according to the level of immune activation in infected humanized mice. The predictive value of EV-associated HIV-1 RNA for immune dysfunction are presently under investigation in a cohort of PLWH. This study shows that characterizing how viral RNA is presented in the plasma, associated with large or small EVs, could provide information overlooked by the standard plasma viral load measurement. Immunotherapy, aiming for a functional cure, gains traction, focusing on controlling viral load by immune cells and restoring the depletion of CD4TL. Notably, the interaction between HIV-1 gp120 and DCIR presents a potential therapeutic target, given its role in viral attachment to DCs and CD4TL, pivotal in AIDS pathogenesis as well as in negative regulators of innate and acquired immune response. Blocking DCIR may restore immune response by inhibiting viral entry and limiting CD4TL depletion and CD8TL exhaustion.

## 4. Methods

### 4.1 Antibodies

For the enzyme-linked immunosorbent assay (ELISA) of HIV-p24, the coating was done using an anti-p24 antibody (hybridoma 183-H12), and detection was done using biotin-conjugated p24 antibody (hybridoma 31-90-25), both antibodies being generous gifts from Dr. Michel Tremblay (Université Laval, QC, Canada). The anti-CD3 (OKT3) and anti-CD28 (9–3) used for spleen cell reactivation were also gifts from Dr. Michel Tremblay. The mouse anti-human CD45-PerCP-Cy5.5 (clone 2D1), CD3-FITC (clone SK7), CD33-PE-Cy7 (clone WM-53), CD8-APC (clone HIT8a), HLA-DR-PE-Cy7 (clone L243) and CD19-APC-eFluor780 (clone SJ25C1) were purchased from Invitrogen. The mouse anti-human PD-1-PE (eBioJ105) and the mouse anti-human CD4-PE (clone RPA-T4) were purchased from BD Biosciences. The mouse anti-human DCIR-Alexa Fluor 405 (216110) was purchased from R&D Systems.

### 4.2 Co-production of virus with EVs mock or EV enriched in miR-155

Two distinct preparation of virus were produced in human embryonic kidney 293T cells (HEK293T) by transient transfection with the calcium/phosphate method, as described previously (111). To produce HIV-1 particles along with miR-155-rich EVs, 2×10^6^ HEK293T cells were transfected simultaneously with 20µg of a pNL4.3Balenv plasmid (provided by R. Pomerantz, Thomas Jefferson University, Philadelphia, PA) and 10 µg of a pMIG-155 plasmid (Addgene cat#26529). To produce HIV-1 particles along with mock EVs, 2×10^6^ HEK293T cells were transfected simultaneously with 20µg of a pNL4.3Balenv plasmid and 10 µg of a pMIGw plasmid (Addgene cat#12282). Cells (2×10^6^ cells per T75 flask) were washed 16 hours after transfection and kept in 10 mL of culture medium for an additional 48 hours. Cell-free supernatant was filtered at 0.22 µm, then centrifuged at 100,000 × g (acceleration = 9; deceleration = 0) for 45 min in an Optima L-90K Beckman Coulter centrifuge with a 70 Ti rotor to pellet virus and EVs. The pellet was resuspended in 150µL of 0.20µm filtered PBS per 10mL of centrifuged culture medium. The viruses were titrated for all stock preparations and experiments using an in-house ELISA for the viral protein p24gag. Both viral preparations containing EV-miR-155 or not were further separated by a velocity gradient as previously shown (81). The gradient was centrifuged at 180,000 x *g* for 50 minutes with slow acceleration and no brakes in a StepSaver 65V13 vertical rotor. MiR-155 was quantified in the viral preparations and the velocity gradient fractions by RT-PCR. The Ct values are presented for the miRNA quantification in the first viral preparations because the standard curve for absolute quantification was not developed at that time. For the second series of experiments, the absolute quantification of miR-155 in the viral preparations was measured by RT-qPCR with a standard curve as described previously (78).

### 4.3 Virus infectivity assay

To titrate virus stock infectivity, we used the TZM-BL indicator cell line which carries a stably integrated luciferase reporter gene under the control of the HIV-1 regulatory element (LTR). In contact with the produced virus, we measured the replication level of a competent infectious virus using luciferase fluorescence detection. Briefly, serial four-fold dilutions of virus stock were titrated in a 96-well plate. Cells were added (10,000 cells / well) and incubated for 48 hours at 37 °C, 5 % CO_2_. Then, cells were lysate with luciferase lysis buffer (25 mM Tris phosphate, pH 7.8, 2 mM dithiothreitol, 1 % Triton X-100 and 10 % glycerol) and kept frozen at −20°C. Twenty-four hours after lysis, an aliquot of cell extract was mixed with luciferin buffer (20 mM tricine, 1.07 mM magnesium carbonate hydroxide pentahydrate, 2.67 mM magnesium sulphate, 100 µM EDTA, 220 µM Coenzyme A, 4.70 µM D-Luciferin potassium salt, 530 µM ATP and 33.3mM dithiothreitol) and luciferase reaction was measured using a Varioskan luminometer (ThermoFisher). The TCID_50_ (50 % Tissue culture infectious dose) was finally calculated by the Spearman-Karber method (112).

### 4.4 NSG-mice model

Female NSG mice were bred at the CRCHU de Québec – Université Laval under the Animal Ethics Committee of Laval University approval and the Canadian Council on Animal Care (CCAC) ethical guidelines. CD34+ hematopoietic stem cells (StemExpress) were incubated at a concentration of 4.0 × 10^5^ cells/mL in StemSpan SFEM medium supplemented with hSCF (100 ng/mL), hflt3 (100 ng/mL), hTPO (50 ng/mL), UM171 (30 nM) (StemCell), plasmocin (5 µg/mL) and primocin (100 µg/mL) (Invivogen). Before cell transplantation, mice were irradiated at 200 cGy using a Gammacell-40 Exactor (Best Theratronics). The day after, 2.5 × 10^5^ cells/mouse were transplanted by intravenous injection in the mouse’s tail. For the first experiment, TCID50 of NL4.3Balenv + EV-Mock (equivalent to 120 ng of p24) or NL4.3Balenv + EV-miR-155 (equivalent to 90ng of p24) was intra-vaginally administrated. For the second experiment, maraviroc (5 mM) or DCIR inhibitor (10 mM) diluted in PBS/2.2 % hydroxyethylcellulose gel was applied intra-vaginally 60 minutes before infection (93). The DCIR inhibitor is also known as 1-benzofuran-2-yl(phenyl)methanone (Chembridge) (29). Non-infected (control) mice received PBS/2.2 % hydroxyethylcellulose. Mice were infected, under anesthesia, by an intra-vaginal injection with the equivalent of 100 ng of p24 of NL4.3Balenv + EV-miR-155.

### 4.5 Blood immune cells isolation

A submandibular puncture was performed to collect blood samples from mice. A euthanasia, blood was collected by a cardiac puncture. Then, mice were under perfusion with HBSS 1X to rinse the remaining blood out of the organs and tissues. Peripheral blood cells were obtained after erythrocytes lysed for 10 minutes at 37°C in an ammonium chloride solution. Cells were washed and resuspended with HBSS 1X. Cells were counted in trypan blue to assess viability with a Cellometer (Nexcelcom Bioscience).

### 4.6 Lungs immune cells isolation

Collected lungs were preserved in RPMI 1640 1X with penicillin (100 U/mL), streptomycin (100 µg/mL) and HEPES (10 mM) until cell purification. Lungs were processed with a gentleMACS dissociator (Miltenyi Biotec Inc.) in HBSS 1X supplemented with Liberase DL (25 µg/mL), DNase I (20 µg/mL) and Elastase (0.0125 U/mL). The dissociated lungs were incubated at 37°C for 30 minutes and processed for a second time with the gentleMACS. Cells were separated from the remaining tissues and extracellular matrix by filtration through a 40 µm-pore size nylon cell strainer (Corning). Lungs cells were centrifuged at 800 x *g* for 30 minutes in the Cytiva Percoll centrifugation media to remove fat and extracellular matrix (113). Erythrocytes were lysed for 5 minutes at 37 °C in an ammonium chloride solution. The remaining cells were washed and diluted in HBSS 1X. Cells were counted in trypan blue to assess viability with a Cellometer (Nexcelcom Bioscience).

### 4.7 Spleen immune cells isolation

Collected spleens were preserved in RPMI 1640 1X with penicillin (100 U/mL), streptomycin (100 µg/mL) and HEPES (10 mM) until cell purification. Cells from the spleen of euthanized mice were obtained using a gentleMACS dissociator in HBSS 1X with fetal bovine serum (2 %) and EDTA (2 mM). The dissociated spleens were processed through a 40 µm-pore size nylon cell strainer (114). Erythrocytes were lysed for 5 minutes at 37 °C in an ammonium chloride solution. The remaining cells were washed and diluted in HBSS 1X. Cells were counted in trypan blue to assess viability with a Cellometer.

### 4.8 Liver immune cells isolation

Collected livers were preserved in RPMI 1640 1X with penicillin (100 U/mL), streptomycin (100 µg/mL) and HEPES (10 mM) until cell purification. Cells were obtained using a gentleMACS dissociator and processed through a 100 µm-pore size nylon cell strainer. Then, PBMCs were isolated through a lymphocyte separation medium. Erythrocytes were lysed for 10 minutes at 37°C in an ammonium chloride solution. Cells were washed and resuspended with PBS. Then, cells were counted in trypan blue to assess viability with a Cellometer.

### 4.9 Bone marrow immune cells isolation

Femurs were cleaned of all muscles and cartilage. The bone marrow was extracted from the femur by a quick centrifugation at 10,000 x *g* (115). Erythrocytes were lysed for 5 minutes at 37 °C in an ammonium chloride solution. The remaining cells were washed and diluted in HBSS 1X. Cells were counted in trypan blue to assess viability with a Cellometer.

### 4.10 Cells flow cytometry

Cells were incubated with FVS450 in HBSS 1X to allow the selection of viable cells. Non-specific binding of antibodies to cells was prevented by incubation with HBSS 1X with fetal bovine serum (1 %), mice serum (1 %) and Human BD Fc Block. Cells were stained with fluorochrome-conjugated antibodies for 30 minutes at room temperature, washed with PBS, and fixed with paraformaldehyde (PFA) 2 % final. Cells were kept at 4 °C until flow cytometer acquisition with a BD FACSCanto II (BD Biosciences).

### 4.11 Spleen cells reactivation

Fresh erythrocyte-free splenic suspensions were suspended in RPMI 10 % ultracentrifuged FBS, 0.1 % primocin (50 mg/mL) and IL-2 (30 U/mL) at 1 × 10^6^ cells/mL. For reactivation, 1.5 × 10^6^ cells were cultured with anti-CD3 (OKT3, 1 µg/mL) / anti-CD28 (9-3, 1 µg/mL) cocktail (activated condition) for nine days. At days 3 and 6, 700 µL of supernatant was removed and replaced with fresh complete culture medium supplemented with IL-2 (30 U/mL).

### 4.12 Plasma EVs purification

Plasma EVs purification was performed according to a well-established procedure (82, 83, 116). Platelet-free plasma (250 µL) was thawed at room temperature and treated with proteinase K (1.25 mg/mL) for 10 minutes at 37 ℃. Plasma was centrifuged at 17,000 × *g* for 30 minutes to pellet large EVs. The remaining supernatant was mixed with 63 µL ExoQuick (System Biosciences) reagent and incubated at 4 ℃ overnight. A centrifugation at 1,500 × *g* for 30 minutes allowed small EVs precipitation. Both small and large EV pellets were washed with 0.22 µm filtered PBS and resuspended in 250 µL of 0.22 µm filtered PBS.

### 4.13 Hydrodynamic size measurement of EVs

EV samples were analyzed in spectrophotometer cuvettes by dynamic light scattering (DLS) with a Zetasizer Nano ZS (Malvern Instruments). Hydrodynamic diameter measurements were done in duplicate at room temperature. Histograms of EVs size distribution allowed the assessment of purified EVS heterogeneity. The measurements also provided the samples derived count rate. This parameter was a approximate relative concentration of EVs in the sample. It was presented as kilo count per second (kcps), meaning the number of photons scattered by EVs per second.

### 4.14 EV quantification by flow cytometry

Absolute EVs quantification by flow cytometry was performed as previously described (80). EVs were stained with lipophilic carbocyanine DiD dye and CellTrace CFSE or Cell Trace Violet (ThermoFisher Scientific), both at a final concentration of 5 µM. Cell Trace Violet was used for EV quantification before infection (Fig S11). Cell Trace CFSE was used for EV quantification at euthanasia (Fig S12). DiD binds and diffuses in the plasma membrane of EVs. Cell Trace crosses the plasma membrane of EVs and binds to amine to become fluorescent. DiD and CFSE double-positive events were considered EVs. In addition, pools of stained EVs were treated with SDS at a final concentration of 0.025% or 0.125% to validate their resistance to detergent (96). EVs treated with SDS were vortexed for 30 seconds and incubated for 30 minutes at room temperature before cytometer acquisition. To determine the EV concentration in our samples, a fixed quantity, measured before every experiment with a Cellometer, of 15 µm silica beads (Polybead Microspheres, Polysciences) was added to every sample. The acquisition was performed on a modified BDFACS canto II with a photomultiplicator on the forward scatter (FSC) channel to improve nanoparticle detection. The number of acquired beads during the cytometer experiments allowed us to calculate the analyzed volume. Then, we divided the absolute number of EVs acquired by the cytometer by the analyzed volume to determine the EV concentration.

### 4.15 Acetylcholine esterase activity assay

Acetylcholinesterase activity was measured following a procedure described previously (117). Briefly, 100 µL of purified EVs fraction were suspended in 100 µL of 1.25 mM acetylthiocholine in PBS pH 8 mixed with 0.1 mM 5,5-dithio-bis(2-nitrobenzoic acid) in PBS pH 7 in a final volume of 200 µL. Changes in absorption were monitored at 450 nm at room temperature for 10 minutes with a SpectraMax 190 plate reader spectrophotometer (Molecular Devices).

### 4.16 RNA extraction

RNA was extracted from 50 µL of purified EVs (diluted in 3 volumes of Trizol LS (ThermoFisher Scientific)) using the phenol/chloroform method and resuspended in 15 µL of Tris/EDTA buffer (118). RNA concentration was measured using a BioDrop spectrophotometer (Montreal Biotech Inc.). RNA quality was considered suitable when the 260 nm/280 nm absorbance ratio was between 1.5 and 2.

### 4.17 Viral load quantification

RT-PCR was performed on 5 µL of RNA with the Superscript IV reverse transcriptase kit (ThermoFisher Scientific) according to the manufacturer instructions with a final concentration of 10 nM of the HIV-1 LTR specific primer set on a GeneAmp PCR System 9700 (Applied Biosystems) (119). The primers were obtained from Integrated DNA Technologies: forward: 5’-GCCTCAATAAAGCTTGCCTTGA-3’; reverse: 5’-GGCGCCACTGCTAGAGATTTT −3′. The pre-amplification step was carried out with the DreamTaq DNA polymerase (ThermoFisher Scientific). Briefly, 10 µL of cDNA was added to a 40 µL mix that contained DreamTaq polymerase buffer (1 X), MgCl_2_ (3 mM), dNTP (300 µM), bovine serum albumin (1 µg/µL), glycerol (2.5 %) and the same primer set (30 nM). The pre-amplification program consisted of 8 minutes at 95 ℃, then 12 cycles of 1 minute at 95 ℃, 3 minutes at 60 ℃ and 1 minute at 72 ℃. Pre-amplified cDNA was diluted by a 10-fold factor with Tris/EDTA buffer before further experiments. QPCR reactions were performed with the QuantiTech SyBr Green kit (Qiagen) according to the manufacturer’s instructions on a CFX384 Touch Real-Time PCR Detection System (Bio-Rad). The final primer concentration was 250 nM.

### 4.18 Anti-p24 ELISA

The viruses were titrated for all stock preparations and experiments using an in-house ELISA for the viral protein p24gag (120).

### 4.19 Statistical analysis

Normal and lognormal distribution tests were carried out for all data sets. Data sets with a lognormal distribution were transformed to obtain a normal distribution for statistical analysis. Student t-tests were used to calculate the difference between two sample groups. One-way ANOVA was performed to compare more than two sample groups at a unique time point. Two-way ANOVA was performed when data for sample groups at multiple time points were available. For correlation analysis in Tables 1, 2 and S2 to S4, the non-parametric Spearman test was performed. Statistical analyses were carried out using GraphPad Prism software version 10.1.2 with p-values below 0.05 considered statistically significant.

## Acknowledgements

This research was funded through Canadian Institutes of Health Research (CIHR) grants; MOP-188726; MOP-267056 (HIV/AIDS initiative) to C.G., and grants from i) Innovation and Economy Minister from Québec Government and SOVAR, ii) Institution fund from Merck Sharpe & Dohme Corp, and iii) CIHR MOP-391232 to CG and F.B. J.B. and W.B.B. are the recipient of the Desjardins scholarship from the Fondation du CHU de Québec. J.B and W.B.B. are recipients of the recruitment scholarship from the AIDS Research Fund of Université Laval. W.W.B. is the recipient of the leadership and sustainable development scholarship and the Fonds de recherche du Québec – Santé (FRQ-S) doctoral training scholarship. The FRQ-S supports the Centre de recherche du CHU de Québec – Université Laval infrastructure. The authors thank Drs. Martin Pelletier and Stephane Gobeil for access to the qPCR platform and Guylaine Jalbert, Veterinarian Daphnée Veilleux, Anne Bergeron, Eva Bresson and Alma Posvandzic for their essential help with the animals. The following reagent was obtained through the NIH AIDS Reagent Program, Division of AIDS, NIAID, Human Immunodeficiency Virus-1 RNA Quantification Standard (#Cat: 3443), Maraviroc (Selzentry) (#Cat: 11580).

## Author contributions

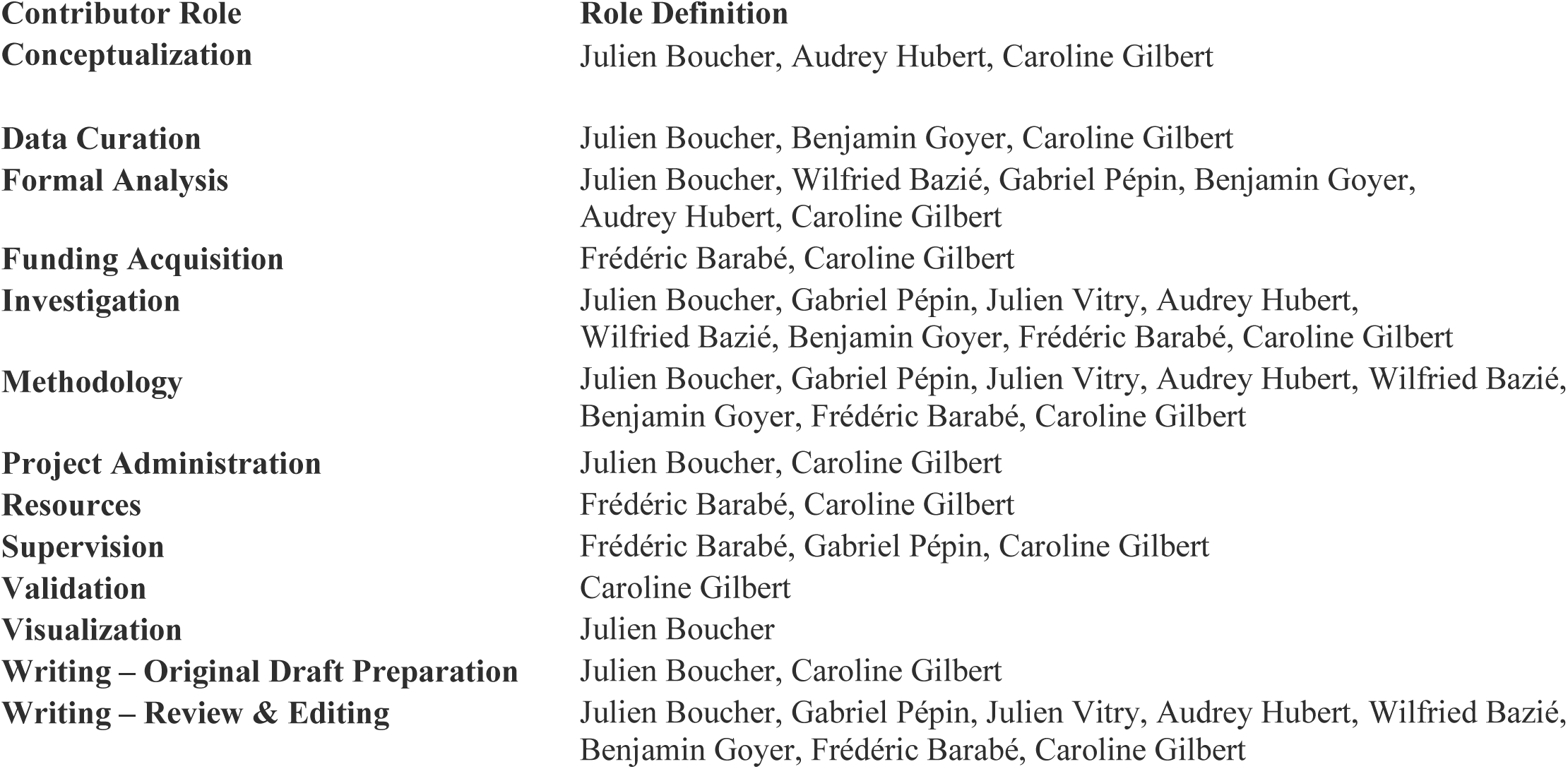

## Institutional review board statement

Animal experiments were previously approved by the “Comité de Protection des Animaux de l’Université Laval (CPAUL)” of the institution.

## Data availability statement

All the data presented in this study are available in both the manuscript and Supporting Information section

## Supporting information

Figure S1. Characterization of the viral preparations.

Figure S2. Optimization of cDNA pre-amplification for EVs viral load measurement.

Figure S3. Modulation of HLA-DR+ and PD-1 in CD4 and CD8 T cells from NL4.3BE/miR-155-infected mice.

Figure S4. Development of an exhaustion phenotype after reactivation of spleen CD8TL.

Figure S5. DCIR expression on purified immune cells from humanized mice.

Figure S6. DCIR inhibition results in a lower plasmatic viral load in humanized mice.

Figure S7. Human immune cells characterization in the spleen, the lungs and the liver of the humanized mice.

Figure S8. Variation of CD8 expression by CD8TL during HIV-1 infection.

Figure S9. Expression of HLA-DR and PD-1 by blood and spleen CD8TL and subsets of CD8^low^ and CD8^high^ T cells.

Figure S10. Gating strategy for EVs quantification by flow cytometry.

Figure S11. Humanized mice plasmatic EV characterization.

Figure S12. Absolute quantification of plasmatic EVs in humanized mice at euthanasia.

Figure S13. Comparison of plasma viral load with EV-associated viral load in humanized mice.

Figure S14. Activation and exhaustion phenotype development in reactivated spleen cells.

Table S1. Effect of pre-amplification on cycle threshold (Ct) values of viral load quantification in plasma EVs.

Table S2. Correlation of mice plasma EVs viral load with biomarkers of disease progression in the spleen.

Table S3. Correlation of mice plasma EVs viral load with biomarkers of disease progression in the lungs.

Table S4. Correlation of mice plasma EVs viral load with biomarkers of disease progression in the liver.

