## Supplementaldata for "Exploring the Relationship Between Extracellular Vesicles, the Dendritic Cell Immunoreceptor and MicroRNA-155 in an In Vivo Model of HIV-1 Infection to Understand the Disease and Develop New Treatments": Supplfigures.pdf

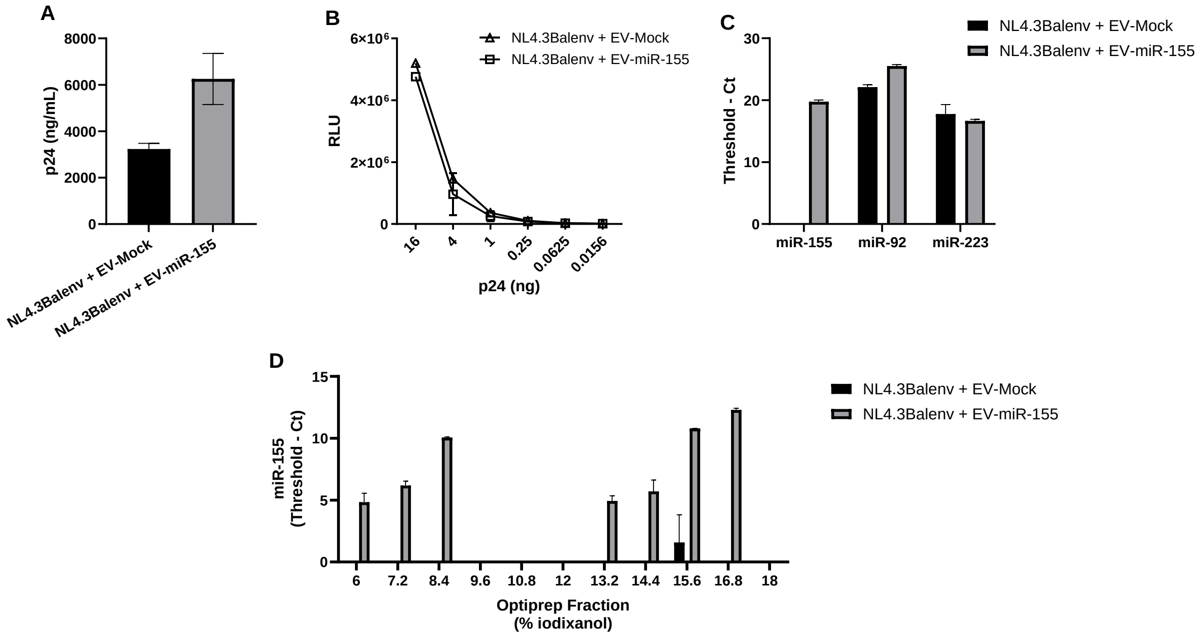


**Figure S1. Characterization of the viral preparations.**
HEK293T cells were co-transfected with an NL4.3Balenv virus-coding plasmid and either a plasmid coding for miR-155 (pMIG-155) or the backbone plasmid (pMIG-w). Cells were washed 16 hours after transfection to remove cell-free plasmids and maintained in culture for 48 hours for virus production. Harvested supernatants were filtered 0.20µm and centrifuged at 100,000 x *g* for an hour to pellet virus and EVs. **A.** Viral capsid protein p24 quantification by ELISA. **B.** Infectivity of the viral preparations measured by luciferase activity in a TZM-bl reporter cell line. **C.** Quantification of miR-155, miR-92 and miR-223 by RT-PCR in the virus and EVs preparations. **D.** The viral preparations were further purified on a velocity gradient of 11 fractions, ranging from 6.0 to 18.0% iodixanol with 1.2% increments. MiR-155 was quantified by RT-PCR in all fractions.

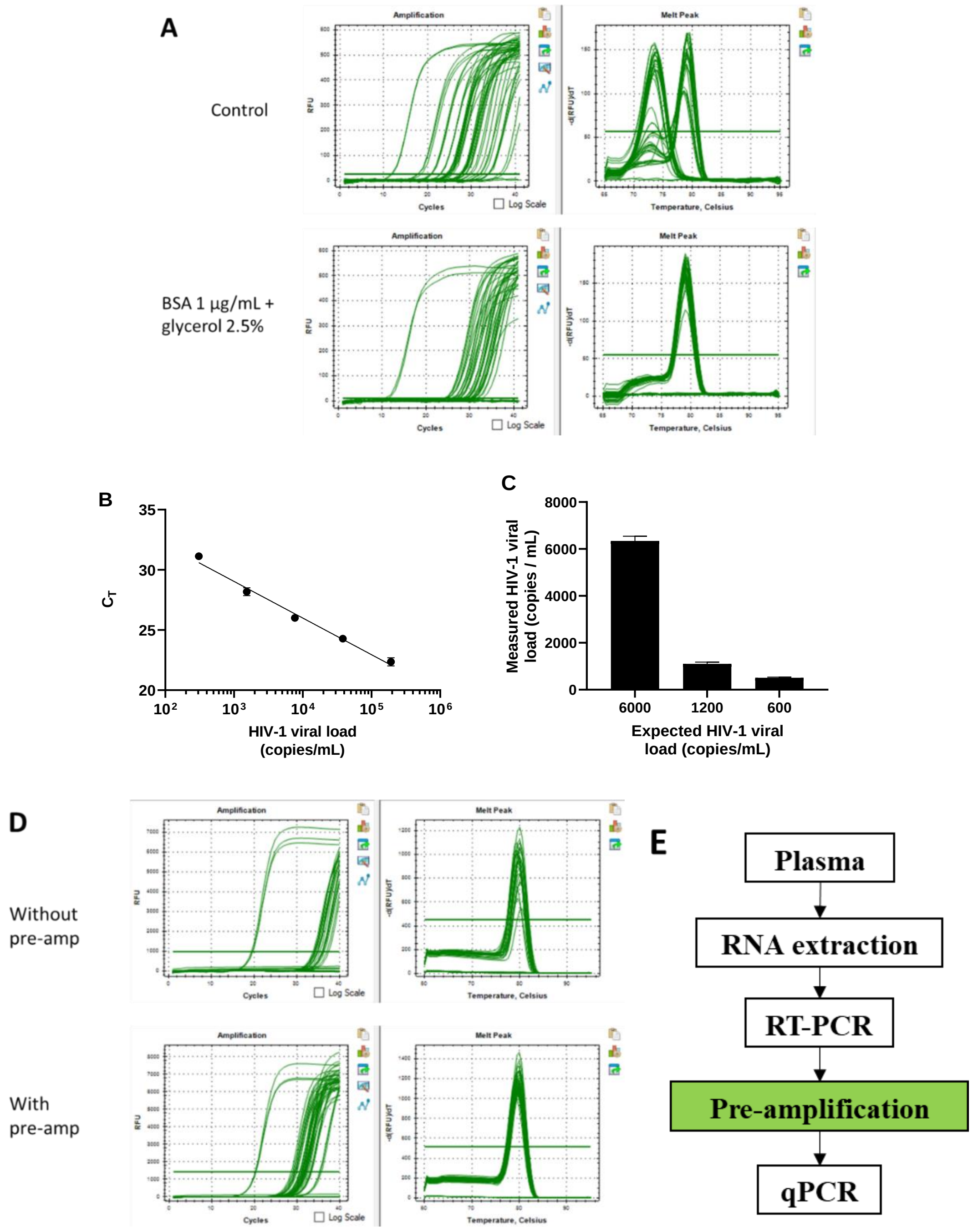


**Figure S2. Optimization of cDNA pre-amplification for EVs viral load measurement.**
**A.** Effect of PCR additives BSA 1 µg/mL and glycerol 2.5 % on the amplification curves and melt curves during qPCR. **B.** Mean of two standard curves produced from two viral preparations of known titer. The standard curve (y = -3.043x + 38.17; $R^2$ = 0.9801) ranges from 192,000 to 300 copies/mL. **C.** NIH HIV RNA standard was diluted to three different dilutions of 6,000, 1,200 and 600 copies/mL. Viral load quantification of the NIH HIV RNA standard with the pre-amplification method. **D.** Effect of pre-amplification on the amplification curves and melt curves during the qPCR experiment of plasma EVs. **E.** Schematic representation of the workflow for the viral load quantification protocol.

**Table S1. Effect of pre-amplification on cycle threshold (Ct) values of viral load quantification in plasma EVs.**

| | no pre-amp | | | pre-amp | | |
|---|---|---|---|---|---|---|
| Sample ID | Ct #1 | Ct #2 | Ct #3 | Ct #1 | Ct #2 | Ct #3 |
| 1 Large EVs | | | | 31.31 | 30.28 | 30.94 |
| 1 Small EVs | | | | 35.04 | 34.93 | |
| 2 Large EVs | | | | | | |
| 2 Small EVs | | | | | | |
| 3 Large EVs | 34.09 | 34.99 | 34.22 | 28.54 | 28.77 | 28.23 |
| 3 Small EVs | | | | | | |
| 4 Large EVs | 36.04 | | | 31.32 | 30.73 | 30.67 |
| 4 Small EVs | 35.95 | 36.03 | | 30.18 | 30.25 | 30.47 |
| 5 Large EVs | | | | 30.93 | 31.10 | 31.40 |
| 5 Small EVs | | | | | | |
| 6 Large EVs | 34.94 | 34.48 | 37.06 | 29.21 | 29.13 | 29.51 |
| 6 Small EVs | | | | 31.78 | 31.68 | 32.60 |

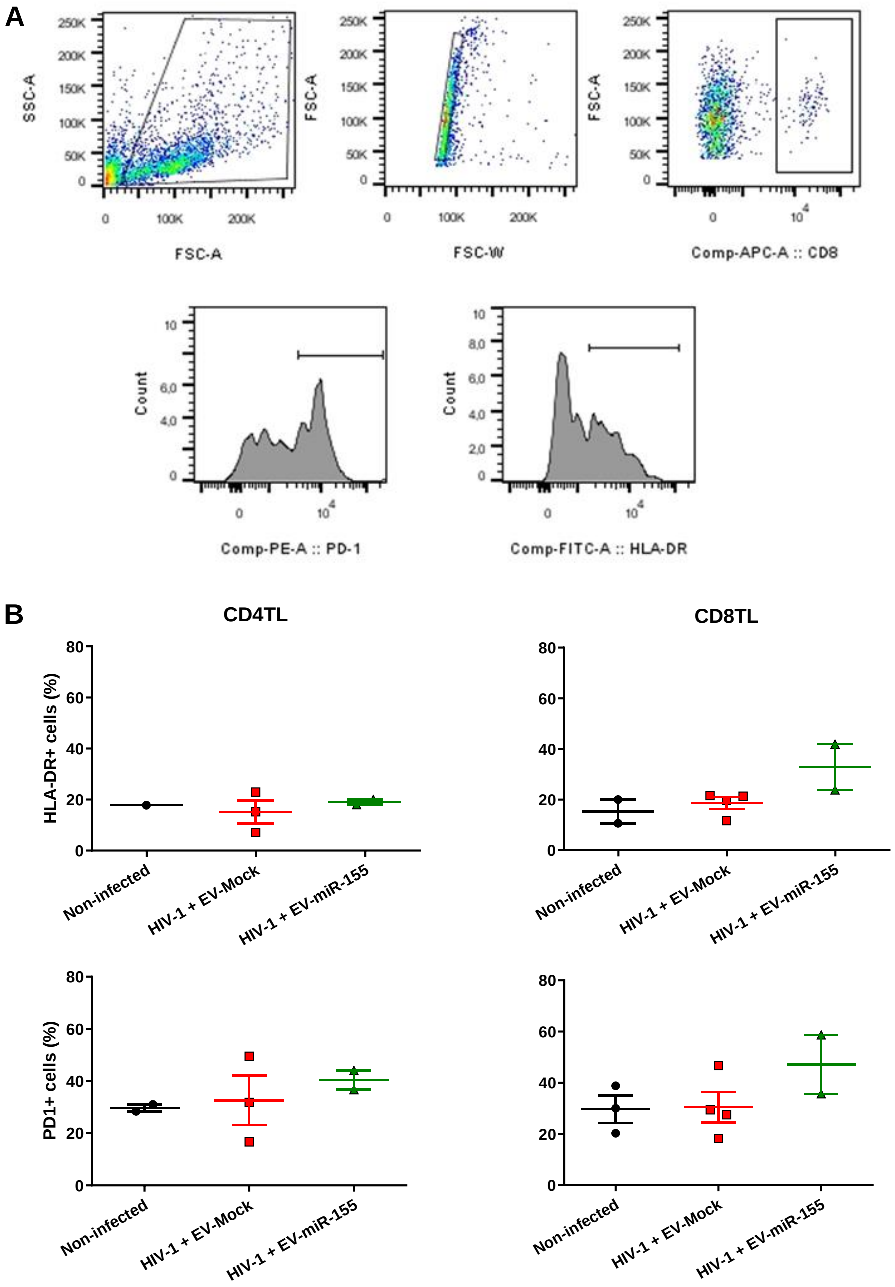


**Figure S3. Modulation of HLA-DR+ and PD-1 in CD4 and CD8 T cells from NL4.3BE/miR-155-infected mice.**

At euthanasia, cells were purified from the blood of control and HIV-1 infected mice and CD4TL and CD8TL were analyzed using flow cytometry. **A**. An example of a gating strategy for PD-1 and HLA-DR+ cells was represented. **B**. Percentages of HLA-DR+ and PD-1+ cells in CD4TL and CD8TL populations were also measured. One-way ANOVA was performed for statistical analysis.

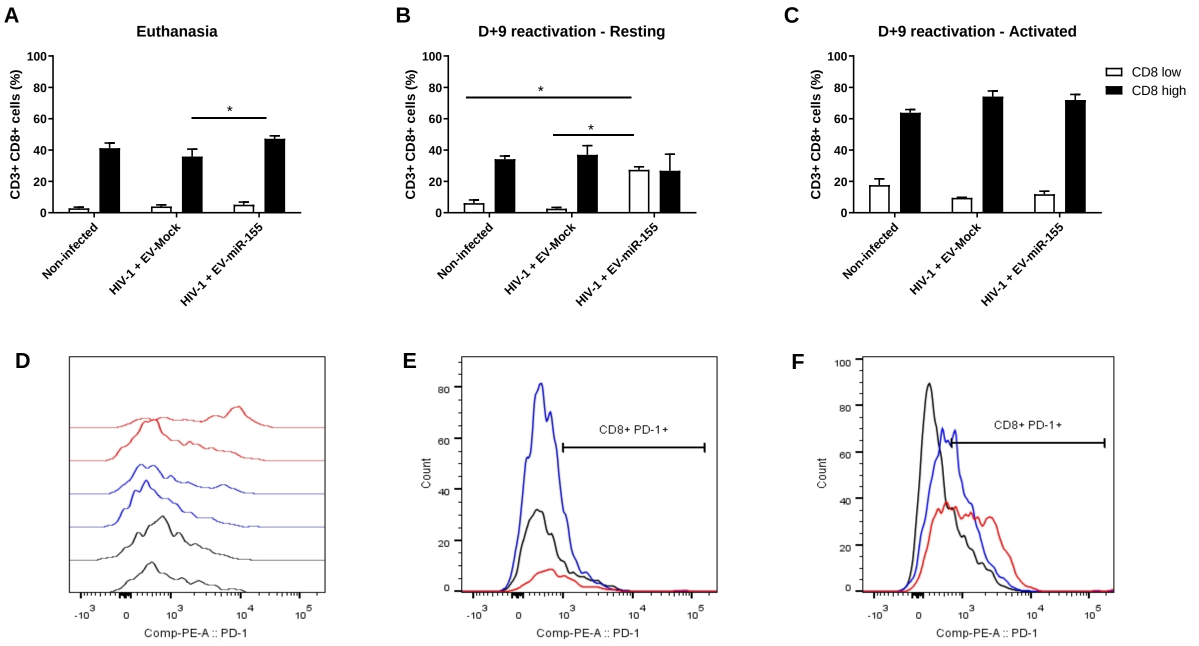


**Figure S4. Development of an exhaustion phenotype after reactivation of spleen CD8TL.** At euthanasia, spleen cells were purified and maintained in culture or activated with an anti-CD3/CD28 cocktail for nine days. Spleen CD3+ CD8+ cells were analyzed using flow cytometry. **A-C**. CD8$^{low}$ and CD8$^{high}$ TL were measured in different groups at euthanasia (A), and after nine days of culture (B) or nine days of activation (C). **D-F**. PD-1+ CD8+ cells were detected. Histograms associated with euthanasia time point (D), nine days resting cells (E) and activated cells (F) were presented. The black line corresponded to the control group, the blue line to the NL4.3BE group and the red line to the NL4.3BE/EV-miR-155 group. A two-way ANOVA analysis with Bonferroni post-test was performed (* $p < 0.05$).

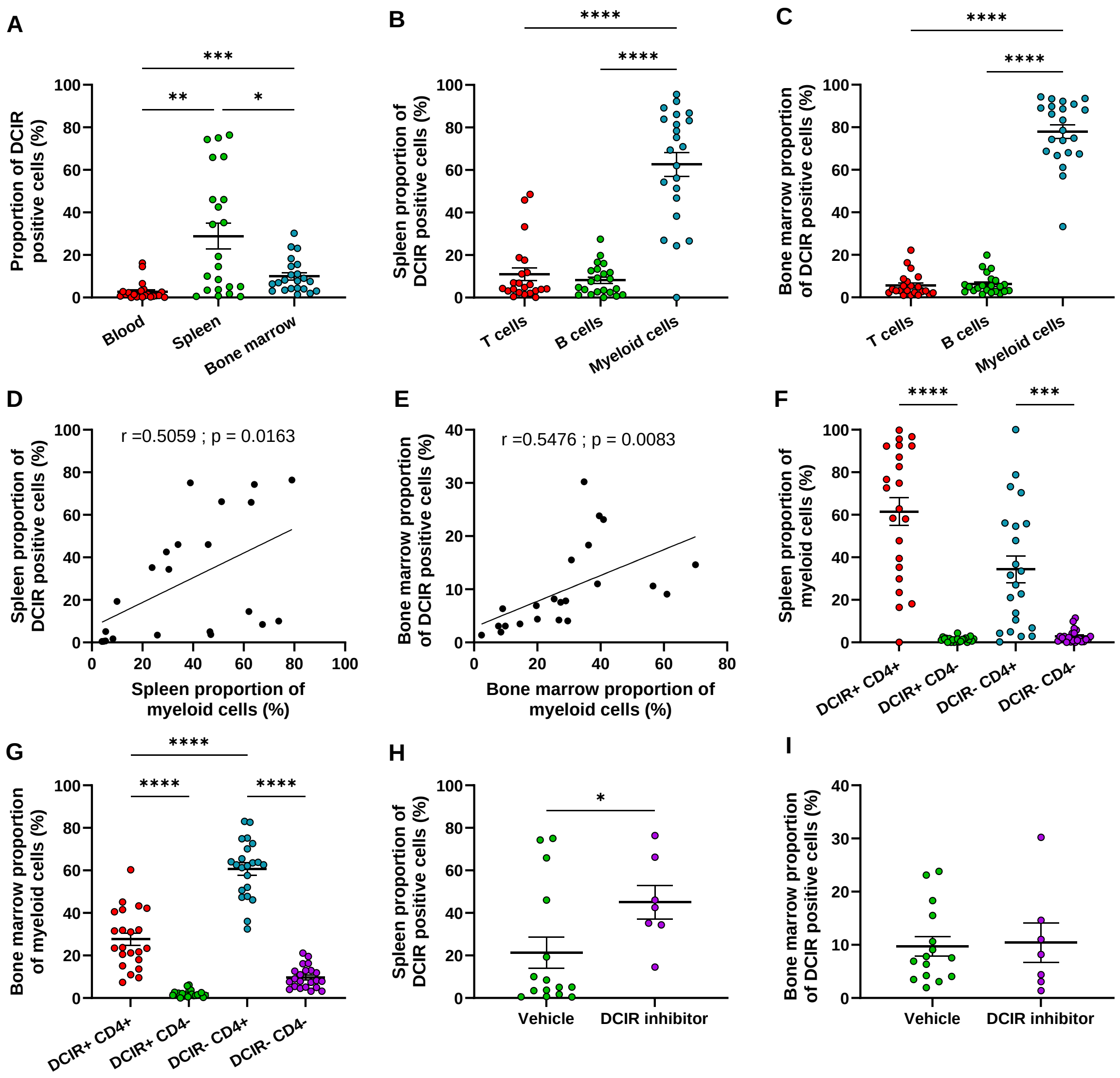


**Figure S5. DCIR expression on purified immune cells from humanized mice.**

**A.** Immune cells were harvested from the blood, the spleen and the bone marrow of humanized mice. Overall DCIR expression was measured on CD45+ cells. **B.** Distribution of DCIR on spleen T cells (CD3+), B cells (CD19+) and myeloid cells (CD33+). **C.** Distribution of DCIR on bone marrow T cells (CD3+), B cells (CD19+) and myeloid cells (CD33+). **D.** Correlation between the proportion of myeloid cells in the spleen and the proportion of DCIR-positive cells. **E.** Correlation between the proportion of myeloid cells in the spleen and the proportion of DCIR-positive cells. **F.** DCIR expression on spleen myeloid cells according to CD4 expression. **G.** DCIR expression on bone marrow myeloid cells according to CD4 expression. One-way ANOVA was performed for statistical analysis (* $p < 0.05$; ** $p < 0.01$; *** $p < 0.001$; **** $p < 0.0001$). **H.** Variation of DCIR expression by spleen cells in response to treatment with the DCIR inhibitor at the moment of infection. **I.** Variation of DCIR expression by bone marrow cells in response to treatment with the DCIR inhibitor at the moment of infection. Non-parametric unpaired t-test was performed for statistical analysis (* $p < 0.05$).

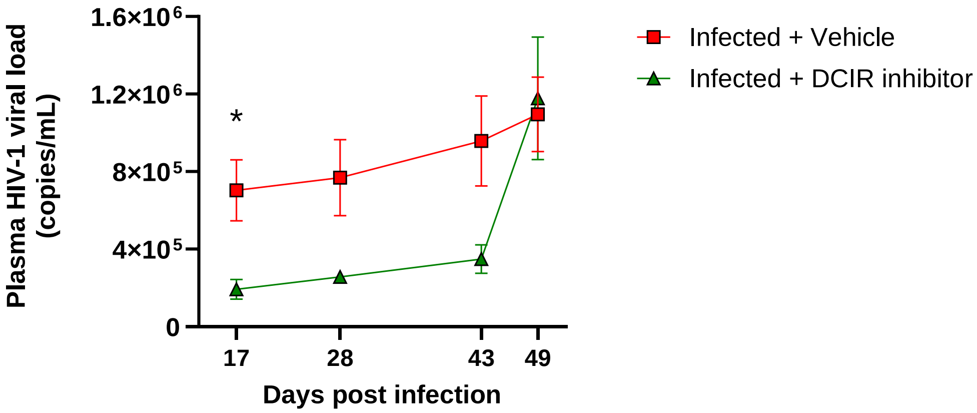


**Figure S6. DCIR inhibition results in a lower plasmatic viral load in humanized mice.** Humanized NSG mice were infected with the NL4.3BE-EV-miR-155 viral preparations. A group of mice were infected with the virus only as the group of reference (n = 7) and another group of mice were inoculated with DCIR inhibitor before infection (n = 4). Plasma viral load was measured by RT-qPCR. An unpaired t-test was performed to compare both groups at each time point independently (* $p < 0.05$).

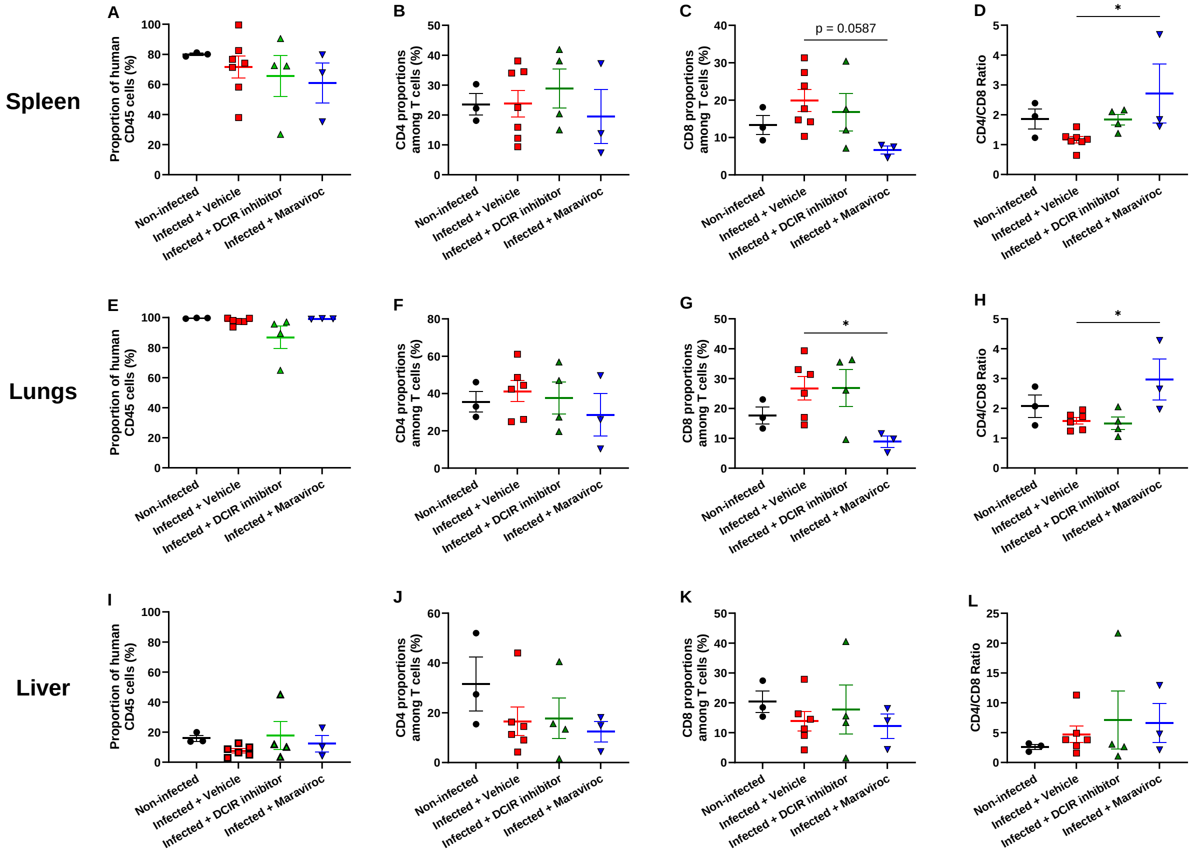


**Figure S7. Human immune cells characterization in the humanized mice's spleen, lungs and liver.**

Spleen, lungs and liver immune cells were purified at euthanasia for flow cytometry analysis. **A-D.** CD45, CD4 and CD8TL proportions were measured among the spleen T cells to calculate the CD4/CD8 ratio. **E-H.** CD45, CD4 and CD8TL proportions were measured among the lung T cells to calculate the CD4/CD8 ratio. **I-L.** CD45, CD4 and CD8TL proportions were measured among the liver T cells to calculate the CD4/CD8 ratio. One-way ANOVA was performed for statistical analysis (* $p < 0.05$).

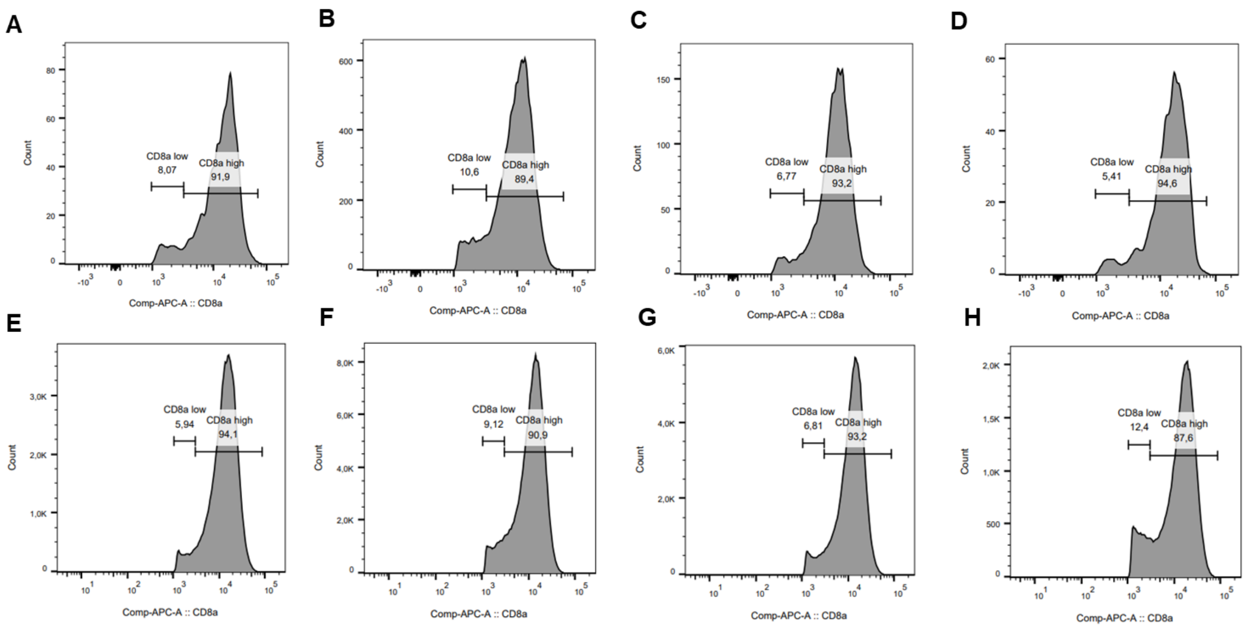


**Figure S8. Variation of CD8 expression by CD8TL during HIV-1 infection.**
Immune cells were purified from the blood (**A-D**) and the spleen (**E-H**) of mice at euthanasia. Total CD8 TL and subsets of CD8$^{low}$ and CD8$^{high}$ TL were analyzed by flow cytometry. **A and E.** Proportions of CD8$^{low}$ and CD8$^{high}$ in non-infected mice. **B and F.** Proportions of CD8$^{low}$ and CD8$^{high}$ in infected mice. **C and G.** Proportions of CD8$^{low}$ and CD8$^{high}$ in infected mice pre-treated with the DCIR inhibitor. **D and H.** Proportions of CD8$^{low}$ and CD8$^{high}$ in infected mice pre-treated with maraviroc.

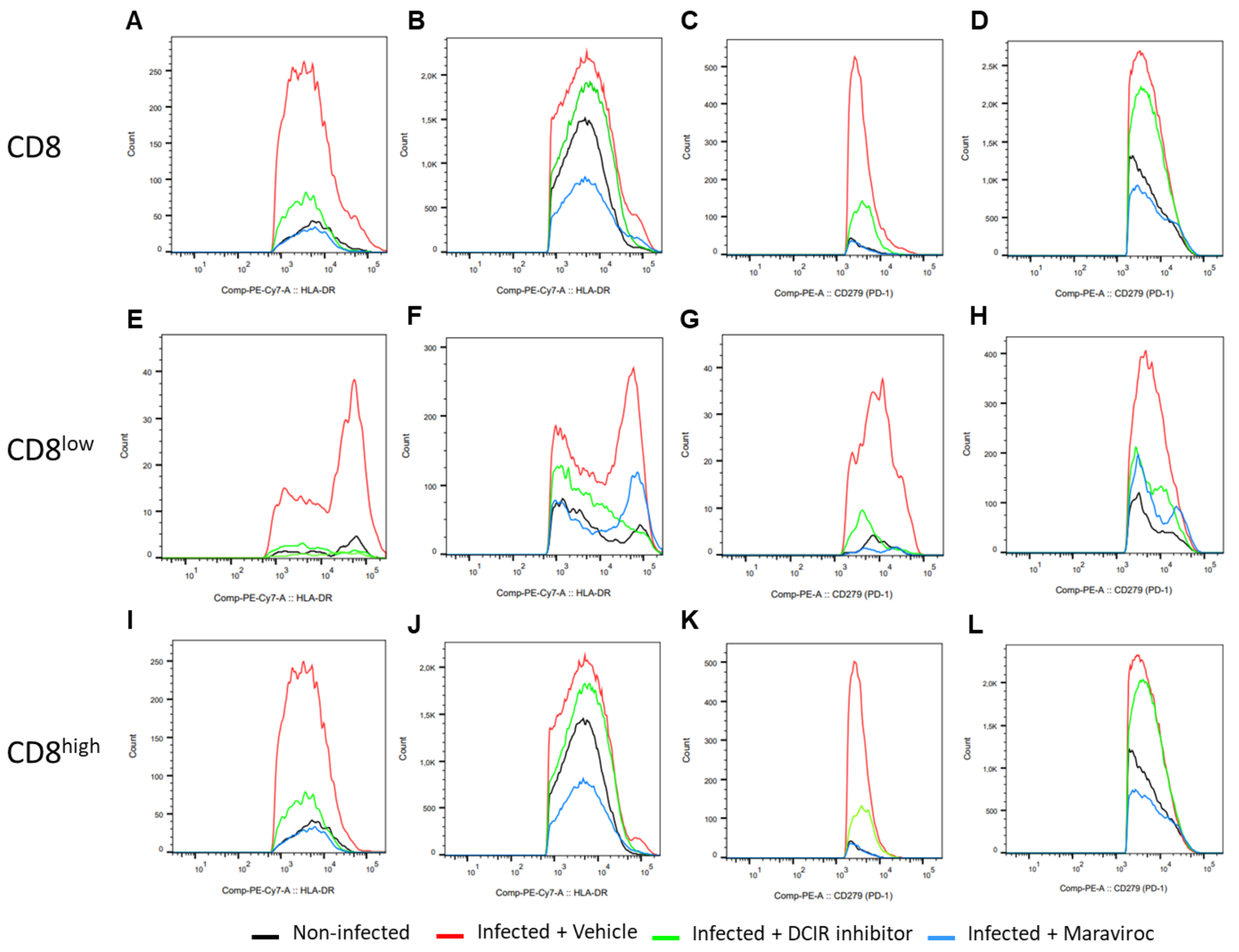


**Figure S9. Expression of HLA-DR and PD-1 by blood and spleen CD8TL and subsets of CD8$^{low}$ and CD8$^{high}$ T cells.**

Immune cells were purified from the blood and the spleen of mice at euthanasia. HLA-DR and PD-1 expression on total CD8TL, CD8$^{low}$ and CD8$^{high}$ TL were characterized by flow cytometry. **A.** Histograms of HLA-DR expression by total blood CD8TL. **B.** Histograms of HLA-DR expression by total spleen CD8TL. **C.** Histograms of PD-1 expression by total blood CD8TL. **D.** Histograms of PD-1 expression by total spleen CD8TL. **E.** Histograms of HLA-DR expression by blood CD8$^{low}$ TL. **F.** Histograms of HLA-DR expression by spleen CD8$^{low}$ TL. **G.** Histograms of PD-1 expression by blood CD8$^{low}$ TL. **H.** Histograms of PD-1 expression by spleen CD8$^{low}$ TL. **I.** Histograms of HLA-DR expression by blood CD8$^{high}$ TL. **J.** Histograms of HLA-DR expression by spleen CD8$^{high}$ TL. **K.** Histograms of PD-1 expression by blood CD8$^{high}$ TL.**L.** Histograms of PD-1 expression by spleen CD8$^{high}$ TL.

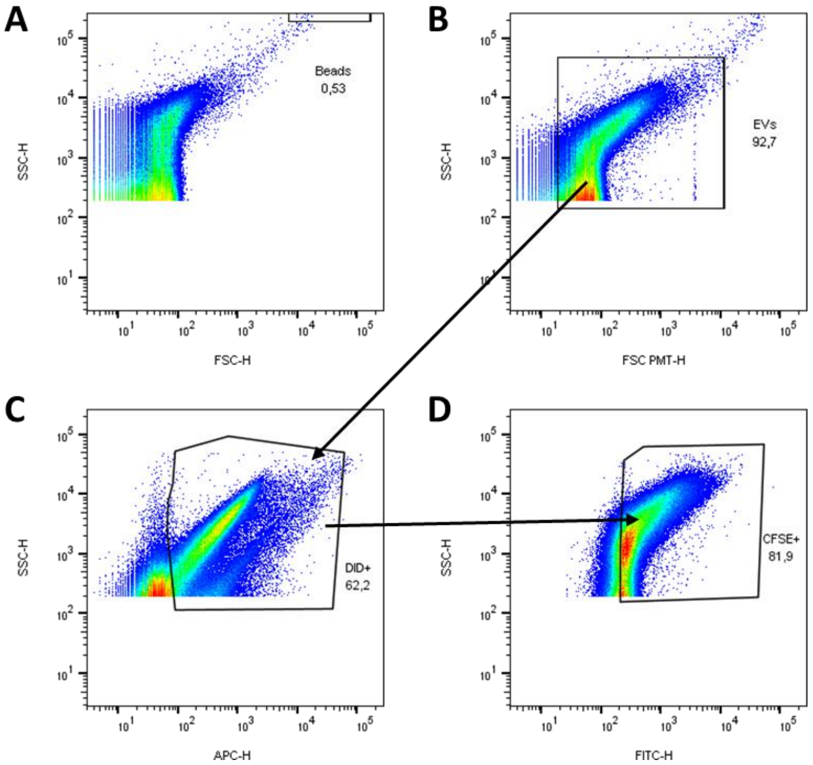


**Figure S10. Gating strategy for EVs quantification by flow cytometry.**
Samples were stained with DiD (5µM), CellTrace CFSE (5µM) and Pluronic 0.02% for 20min at 37°C. Then samples were diluted 1/100 in a final volume of 500µL. 5µL of beads 15 µm (with known concentration) was added and samples were analyzed using flow cytometry. We defined the stopping gate on the beads gate at 1000 events. **A.** Count beads were selected by size (FSC) and granularity (SSC). **B.** Total nanoparticles were selected according to size (FSC-PMT, photomultiplicator) and granularity (SSC). The photomultiplicator allows a better distinction between nanoparticles and extracellular debris. **C.** Among total nanoparticles, only DiD+ (APC) particles were analyzed. DiD is dye-specific for lipid bilayers. DiD- events are excluded because they are considered non-vesicular extracellular material. DiD tends to create false positive events. To exclude false positive events, EVs were also marked with CellTrace CFSE. **D.** CellTrace CFSE+ (FITC) events were selected from the DiD+ gate only. Thus, DiD and CellTrace double-positive events were considered as EVs.

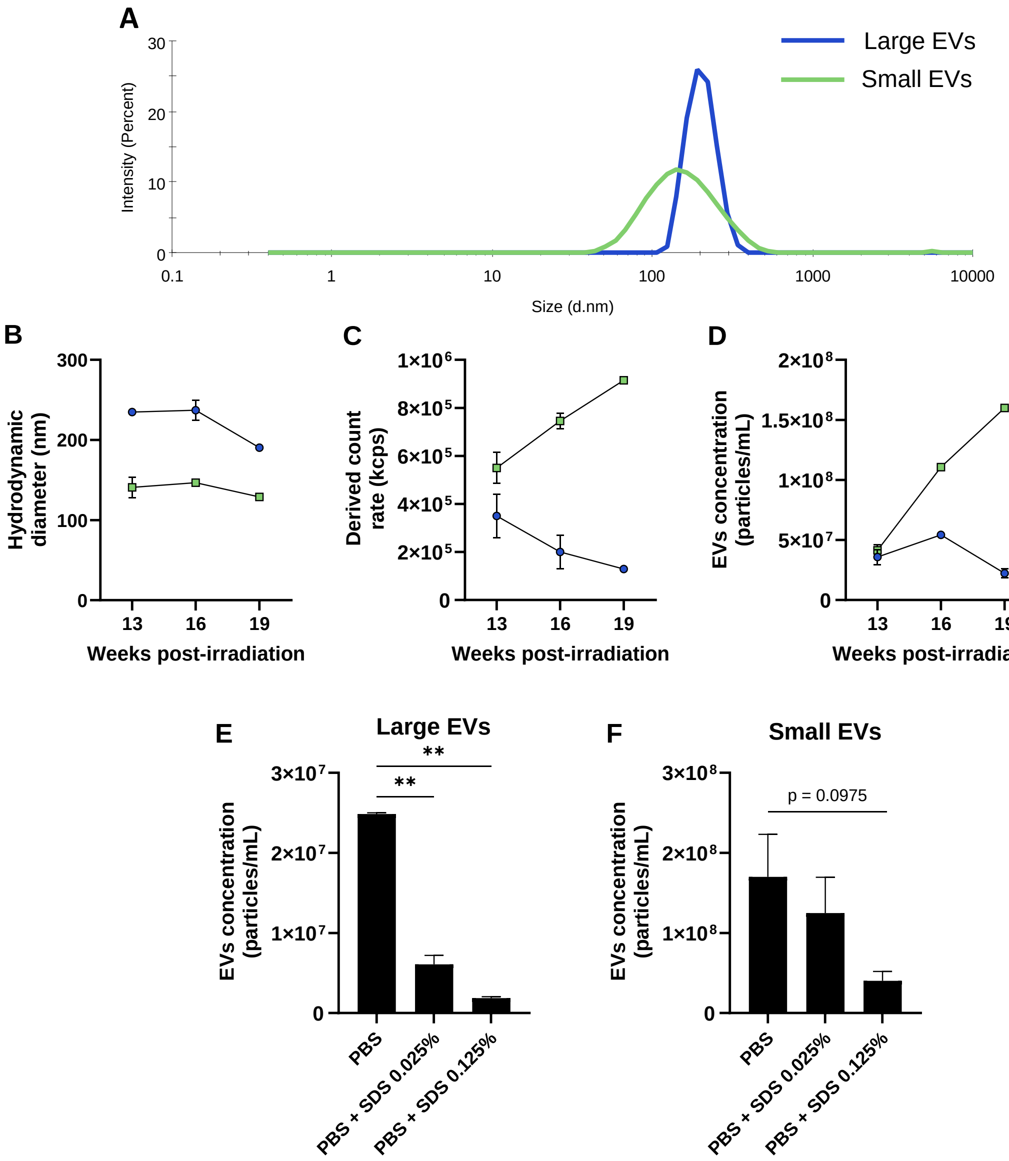


**Figure S11. Humanized mice plasmatic EV characterization.**
After mice humanization, the plasma of every mouse was pooled at each blood sampling (13, 16, and 19 weeks post-irradiation) for EV characterization. **A.** Hydrodynamic diameter of purified EVs measured by dynamic light scattering with Nanosizer (the mean of the three samples is shown). **B.** The average diameter of all measured nanoparticles in the sample. **C.** Derived count rate in kilo count per second (kcps) provided by the Nanosizer analysis. The derived count rate is the number of photons scattered by EVs per second during the sample reading. It allows a relative EVs concentration comparison between samples. **D.** Purified EVs stained with DiD and Cell Trace dyes for absolute quantification by flow cytometry. Pools of **E.** large EVs or **F.** small EVs were treated with final concentrations of 0.025% or 0.125% to assess their resistance to detergent. One-way ANOVA statistic tests were performed for statistical comparisons (** $p < 0.01$).

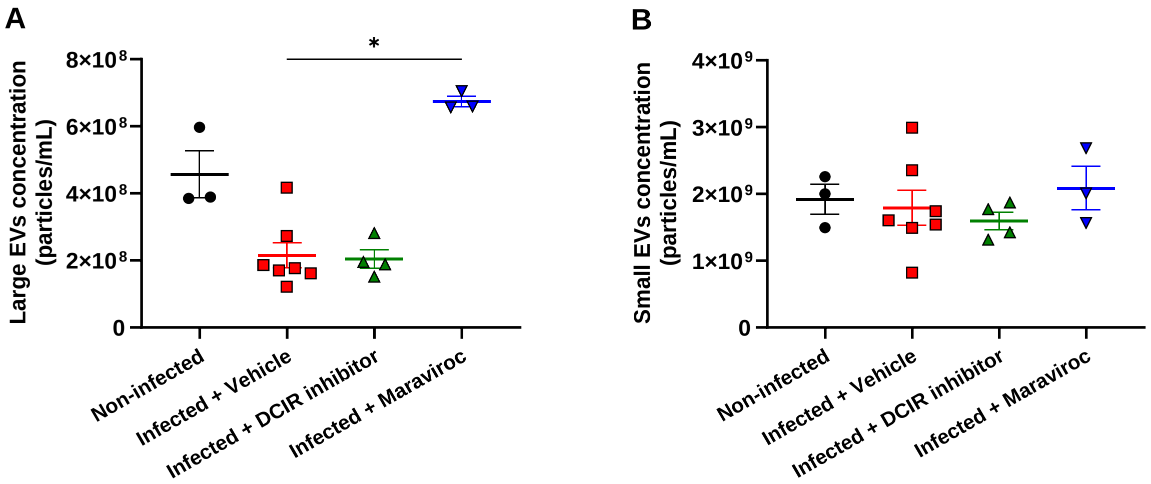


**Figure S12. Absolute quantification of plasmatic EVs in humanized mice at euthanasia.** Absolute quantification by flow cytometry of **A.** large EVs and **B.** small EVs purified from the plasma of humanized mice. A one-way ANOVA statistic test was performed (* $p < 0.05$).

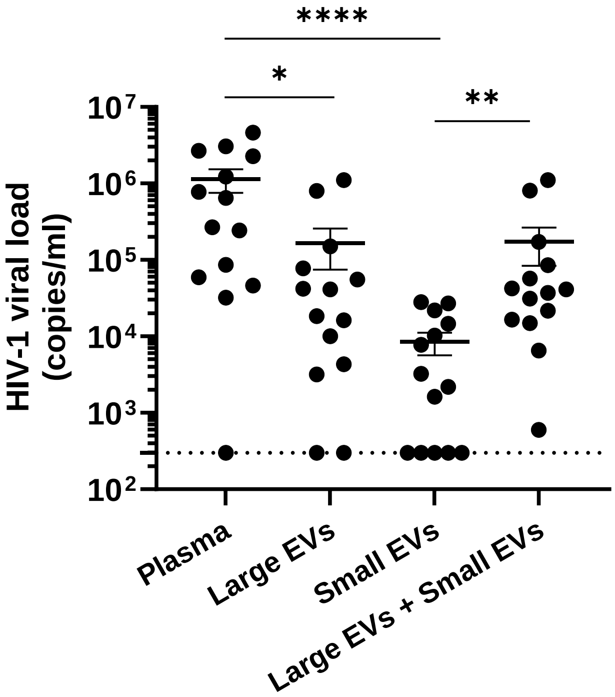


**Figure S13. Comparison of plasma viral load with EV-associated viral load in humanized mice.**
Quantification of viral load in the total plasma and EVs purified from the plasma of mice. A one-way ANOVA statistic test was performed (* $p < 0.05$; ** $p < 0.01$; **** $p < 0.0001$).

**Table S2. Correlation of mice plasma EVs viral load with biomarkers of disease progression in the spleen.**

| | CD4+ T cells count | CD8+ T cells count | CD4/CD8 ratio |
|---|---|---|---|
| Plasma viral load (copies/mL) | r = -0.0339; p = 0.9084 | r = 0.2324; p = 0.4239 | r = -0.3099; p = 0.2809 |
| Large EVs viral load (copies/mL) | r = 0.4260; p = 0.1288 | **r = 0.4828; p = 0.0804** | r = -0.1492; p = 0.6107 |
| Large EVs viral load (copies/EV) | r = 0.4581; p = 0.0995 | **r = 0.5308; p = 0.0508** | r = -0.1555; p = 0.5956 |
| Small EVs viral load (copies/mL) | r = -0.0693; p = 0.8139 | r = -0.1158; p = 0.6934 | r = 0.2206; p = 0.4486 |
| Small EVs viral load (copies/EV) | r = 0.04266; p = 0.8849 | r = 0.1892; p = 0.5172 | r = -0.0247; p = 0.9331 |

**Table S3. Correlation of mice plasma EVs viral load with biomarkers of disease progression in the lungs.**

| | CD4+ T cells count | CD8+ T cells count | CD4/CD8 ratio |
|---|---|---|---|
| Plasma viral load (copies/mL) | r = -0.2836; p = 0.3478 | r = 0.2539; p = 0.4026 | r = -0.3413; p = 0.2538 |
| Large EVs viral load (copies/mL) | r = -0.1199; p = 0.6964 | r = 0.2276; p = 0.4546 | r = -0.2040; p = 0.5059 |
| Large EVs viral load (copies/EV) | r = -0.1214; p = 0.6929 | r = 0.2308; p = 0.4481 | r = -0.2211; p = 0.4678 |
| Small EVs viral load (copies/mL) | r = 0.1988; p = 0.5150 | r = -0.1603; p = 0.6009 | r = -0.1212; p = 0.6933 |
| Small EVs viral load (copies/EV) | r = 0.0346; p = 0.9106 | r = -0.0969; p = 0.7527 | r = -0.1352; p = 0.6597 |

**Table S4. Correlation of mice plasma EVs viral load with biomarkers of disease progression in the liver.**

| | CD4+ T cells count | CD8+ T cells count | CD4/CD8 ratio |
|---|---|---|---|
| Plasma viral load (copies/mL) | r = 0.2786; p = 0.3567 | r = 0.3254; p = 0.2780 | r = -0.2034; p = 0.5051 |
| Large EVs viral load (copies/mL) | r = 0.5204; p = 0.0683 | r = 0.4894; p = 0.0896 | r = -0.1833; p = 0.5489 |
| Large EVs viral load (copies/EV) | r = 0.5415; p = 0.0560 | r = 0.5212; p = 0.0678 | r = -0.1941; p = 0.5252 |
| Small EVs viral load (copies/mL) | r = -0.2861; p = 0.3433 | r = -0.2514; p = 0.4073 | r = 0.1228; p = 0.6894 |
| Small EVs viral load (copies/EV) | r = -0.1723; p = 0.5735 | r = 0.0019; p = 0.9950 | r = -0.1264; p = 0.6808 |

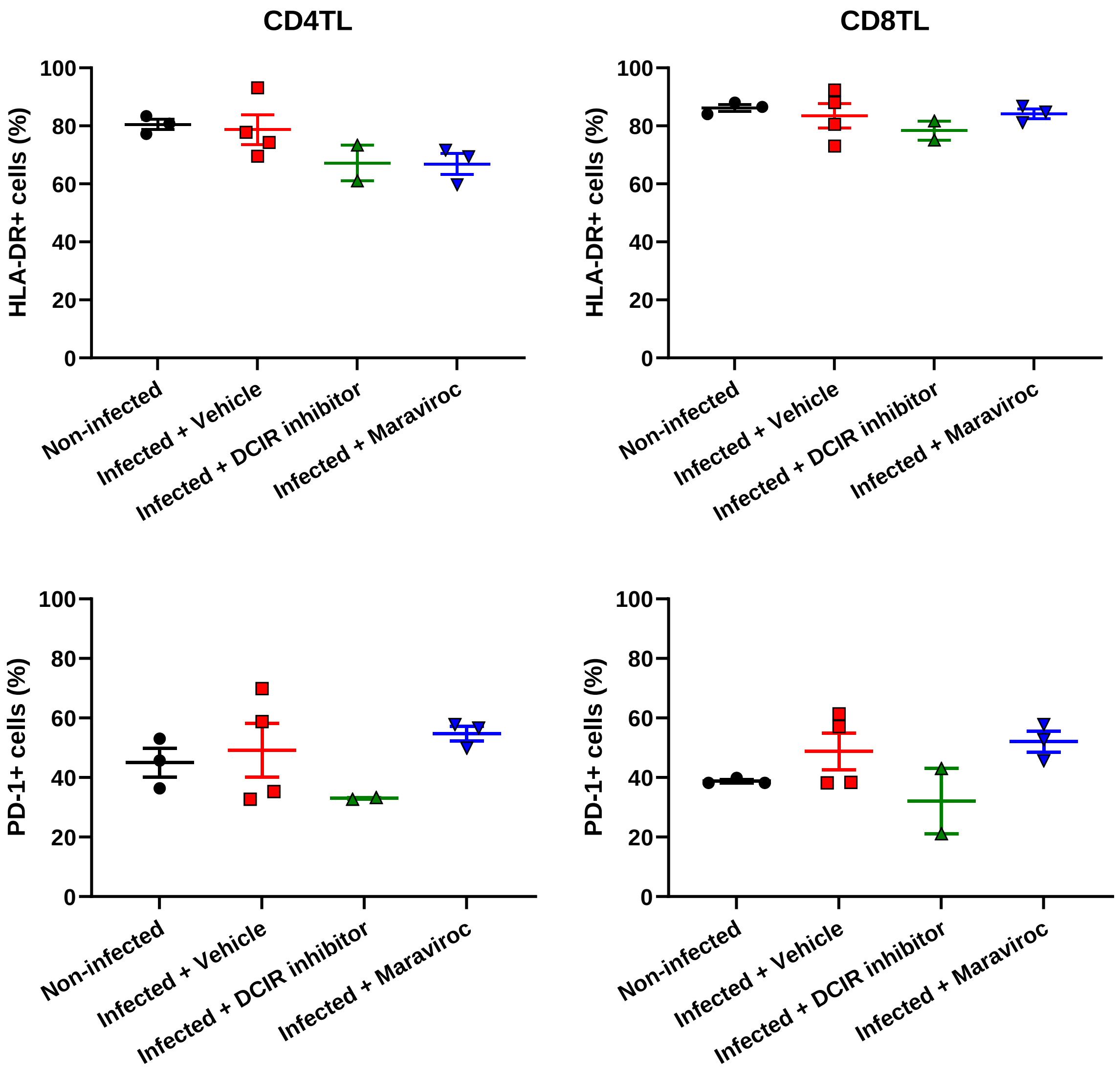


**Figure S14. Activation and exhaustion phenotype development in reactivated spleen cells.** At euthanasia, spleen cells were purified and maintained in culture or activated with an anti-CD3/CD28 cocktail for nine days. Spleen CD3+ CD8+ cells were analyzed using flow cytometry. CD3+ CD8- were considered CD4TL. HLA-DR and PD-1 expression were both measured in CD4 and CD8TL. Percentages of HLA-DR+ and PD-1+ cells in CD4TL and CD8TL populations are shown. One-way ANOVA was performed for statistical analysis.
